# Dual modulation of phase-transitioning licenses the Bicc1 network of ciliopathy proteins to bind specific target mRNAs

**DOI:** 10.1101/2021.10.07.463531

**Authors:** Benjamin Rothé, Simon Fortier, Daniel B. Constam

## Abstract

Perturbations in biomolecular condensates that form by phase-transitioning are linked to a growing number of degenerative diseases. For example, mutations in a multivalent interaction network of the Ankyrin (ANK) and sterile alpha motif (SAM) domain-containing ANKS3 and ANKS6 proteins with the RNA-binding protein Bicaudal-C1 (Bicc1) associate with laterality defects and chronic kidney diseases known as ciliopathies. However, insights into the mechanisms that control RNA condensation in ribonucleoprotein particles (RNPs) are scarce. Here, we asked whether heterooligomerization modulates Bicc1 binding to RNA. Reconstitution assays *in vitro* and live imaging *in vivo* show that a K homology (KH) repeat of Bicc1 self-interacts and synergizes with SAM domain self-polymerization independently of RNA to concentrate bound mRNAs in gel-like granules that can split or fuse with each other. Importantly, emulsification of Bicc1 by ANKS3 inhibited binding to target mRNAs, whereas condensation by ANKS6 co-recruitment increased it by liberating the KH domains from ANKS3. Our findings suggest that the perturbation of Bicc1-Anks3-Anks6 RNP dynamics is a likely cause of associated ciliopathies.

## INTRODUCTION

Liquid-liquid or liquid-gel phase separations can segregate molecules from their solute into supramolecular polymer condensates of variable size and viscosity such as stress granules (SGs), P-bodies (PBs), centrosomes and the nucleolus (Boeynaems et al., 2018; Riggs et al., 2020). PBs and SGs accrue proteins that participate in RNA metabolism and translational control, including Bicc1 (Estrada Mallarino et al., 2020; Maisonneuve et al., 2009; Rothé et al., 2015; Tran et al., 2010; Youn et al., 2019, 2018). Condensation of these membraneless organelles is nucleated by one or several scaffold proteins with intrinsically disordered regions or RNA-binding domains that cooperate to assemble multivalent protein-protein and protein-RNA interaction networks. In addition, phase separation can be modulated by intermolecular RNA-RNA contacts, and by changes in pH, salinity and temperature to allow rapid integration of environmental cues, and in a reversible manner (Alberti et al., 2019; Boeynaems et al., 2018; Van Treeck and Parker, 2018). Perturbations which instead favor irreversible aggregation are implicated in a growing number of diseases (Alberti and Hyman, 2021). Therefore, considerable efforts are needed to delineate the multivalent molecular interactions and their hierarchy that govern the dynamics and the function of such RNA-protein networks.

Bicc1 consists of a tandem repeat of three KH and two KH-like (KHL) domains at the N-terminus, separated from a self-polymerizing C-terminal SAM domain by a serine- and glycine-rich intervening sequence (IVS). Analysis of the protein interactomes of Bicc1 and its homolog in *Drosophila* confirmed a physical connection with core components of PBs, including the carbon catabolite repressor 4–negative on TATA (Ccr4-Not) complex, and with factors which promote SG formation in stressed cells (Chicoine et al., 2007; Kugler et al., 2009; Leal-Esteban et al., 2018; Youn et al., 2018). Ccr4-Not mediates translational repression and decay of miRNA-induced silencing complexes (miRISC) (Collart and Panasenko, 2012) and PB condensation by the RNAi machinery (Eulalio et al., 2007). Alternatively, Ccr4-Not can be recruited to mRNA by specific proteins (Temme et al., 2014). Consistent with a role in activating Ccr4-Not, *Drosophila* Bicc1 binds the Cnot3/5 subunit and promotes deadenylation of its own mRNA during the initial stages of oogenesis (Chicoine et al., 2007). In addition, loss-of-function studies revealed that Cnot3 and Bicc1 accelerate the decay of *Dand5* mRNA to specify the left-right asymmetry of visceral organs in vertebrates (Maerker et al., 2021; Minegishi et al., 2021). Regulated Dand5 mRNA decay depends on the proximal 3’-UTR, which is recognized by the KH_1_ and KH_2_ domains of Bicc1 via a bipartite GAC motif in the conserved GACGUGAC sequence, and on flow stimulation of polycystin-2 in primary cilia on the left side of the node (Minegishi et al., 2021). However, how polycystin-2 activates Bicc1 is unknown. *Bicc1* mutations also provoke the development of fluid-filled cysts in kidneys, liver and pancreas that are reminiscent of autosomal dominant polycystic kidney disease (ADPKD) (Cogswell et al., 2003; Kraus et al., 2012). In polycystin mutant kidneys, epithelial cell proliferation and fluid secretion are stimulated by excessive accumulation of cAMP (Wallace, 2011). Cyclic AMP also increases in Bicc1 mutant kidneys, possibly due to de-repression of adenylate cyclase 6 (Adcy6) mRNA translation (Piazzon et al., 2012). Bicc1-mediated silencing requires the KH domains to recruit *Adcy6* mRNA, whereas self-polymerization of the SAM domain slows Bicc1 turnover and concentrates bound mRNA in cytoplasmic granules (Piazzon et al., 2012; Rothé et al., 2015). *Adcy6* transcripts can be translationally silenced by Bicc1 also in the absence of flow and independently of deadenylation and decay (Piazzon et al., 2012). This raises the question of how the fate of Bicc1-bound mRNAs might be regulated by transcript-specific interacting factors.

The sterile alpha motif (SAM) consists of a bundle of five α-helices that form a small globular domain in over 1,300 eukaryotic proteins, often to mediate their head-to-tail polymerization in helical arrangements via their mid-loop (ML) and end-helix (EH) surfaces (Bienz, 2020; Kim et al., 2001). Bicc1 is the only protein known to combine a SAM domain with RNA-binding KH domains (Letunic et al., 2021). In crystals, the SAM domain forms a flexible polymer scaffold that is predicted to arrange the KH repeat and a serine- and glycine-rich intervening sequence (IVS) at its periphery (Rothé et al., 2015, 2018). However, a role for RNA or other clients of KH domains in Bicc1 phase separation remains to be tested. Known Bicc1-interacting factors include ANKS3 and ANKS6 (Bakey et al., 2015; Nakajima et al., 2018; Rothé et al., 2018; Stagner et al., 2009; Yakulov et al., 2015), which are mutated in patients and animal models with left-right patterning defects or kidney cysts, or both (Bakey et al., 2015; Brown et al., 2005; Czarnecki et al., 2015; Hoff et al., 2013; Shamseldin et al., 2016; Stagner et al., 2009; Taskiran et al., 2014). Bicc1 and ANKS3 bind each other via SAM:SAM interactions and through additional contacts mediated by their KH and Ank repeats, and by their respective IVS (Rothé et al., 2018). The region of Bicc1 accommodated by the rigid Ank platform is unknown. However, reconstitution experiments in heterologous cells revealed that ANKS3 hinders the elongation of Bicc1 polymers due to its bulky C-terminus and thereby disperses cytoplasmic Bicc1 granules. Alternatively, ANKS3 can recruit ANKS6 (Leettola et al., 2014; Rothé et al., 2018). By sequestering the ANKS3 SAM domains with high affinity, ANKS6 frees the Bicc1 SAM domain to self-polymerize in large Bicc1-ANKS3-ANKS6 condensates (Leettola et al., 2014; Rothé et al., 2018). Elucidating the relative contributions of individual protein domains to such condensates will be critical for any future therapeutics that may target the underlying multivalent interaction network.

Here, we investigate the roles of specific RNA and protein clients of the KH repeat in Bicc1 phase transitioning, and whether phase separation in turn influences RNA binding. Using *in vitro* reconstitution experiments and live imaging of GFP-tagged Bicc1, we show that the KH repeat contributes to phase separation independently of bound RNA, and that the resulting gel-like condensates show actomyosin-dependent mobility and dynamically exchange associated GFP-Bicc1 transcripts with the soluble phase. In addition, we identify an interaction of ANKS3 with Bicc1 KH domains that inhibits their association with *Dand5*, *Adcy6* and *Bicc1* mRNAs. Conversely, ANKS6 is shown to weaken the interaction of Bicc1 with ANKS3 and to increase RNA binding. Taken together, our data suggest that ANKS3 and ANKS6 act as chaperones of Bicc1 to couple its RNA binding with phase separation.

## RESULTS

### The KH domain repeat of Bicc1 mediates RNA-independent low speed sedimentation in vitro

To assess its solubility, endogenous Bicc1 in cytoplasmic extracts of murine inner medullary collecting duct (IMCD3) cells was subjected to a protocol of differential centrifugation that enriches membraneless organelles (Teixeira et al., 2005; Jain et al., 2016; Namkoong et al., 2018). Western blot analysis showed that 30% of endogenous Bicc1 partitioned to heavy fractions (**Figure 1A-B**). To characterize Bicc1 oligomers in the heavy fractions by fluorescent imaging, we expressed doxycycline-inducible GFP-tagged HA-Bicc1 (GFP-Bicc1) in IMCD3 cells where endogenous *Bicc1* was disrupted by a frame-shifting indel mutation in exon 1 using CRISPR/Cas9 editing to prevent co-polymerization with GFP-Bicc1 (Leal-Esteban et al., 2018). Western blot analysis and live imaging showed that doxycycline induced GFP-Bicc1 up to 6-fold above endogenous levels and formed granules within less than 6 hrs in >95% of cells (**Figure 1C, Figure 1 - figure supplement 1A-B**). Differential centrifugation of cell extracts revealed over 60% of GFP-tagged HA-Bicc1 in the 2000-4000g heavy fractions containing the RNA granule marker CNOT1 (Parker and Sheth, 2007), indicating that overexpression or fusion to GFP further reduces Bicc1 solubility (**Figure 1D**). Nevertheless, even at this elevated expression level, the addition of ≥300 mM NaCl to cell extraction buffer largely abolished GFP-Bicc1 sedimentation, suggesting that it is mediated by stereospecific interactions rather than by non-specific aggregation (**Figure 1 - figure supplement 1C**). Moreover, confocal imaging after embedding in low melting agarose revealed the presence of micrometer-scale GFP fluorescent granules in the 4,000 g fraction of doxycycline-treated but not of uninduced control cells or in the light fraction (**Figure 1 - figure supplement 1D**). To assess whether these Bicc1 sediments correspond to the cytoplasmic granules, we transiently transfected HEK293T cells with full-length HA-Bicc1 or ΔKH or ΔSAM mutants lacking the KH or SAM domains. We chose HEK293T cells because they express no endogenous BICC1 that could interfere with HA-Bicc1. Deletion of the SAM domain reduced HA-Bicc1 partitioning to heavy fractions by less than 10%, whereas deletion of the KH domains shifted it almost entirely to the light fraction (**Figure 1E**). This result was unexpected since previous analysis established that cytoplasmic Bicc1 protein granules can form independently of KH domains by head-to-tail association of the SAM domain (Knight et al., 2011; Maisonneuve et al., 2009; Rothé et al., 2015). To evaluate whether the KH repeat mediates Bicc1 sedimentation in cell lysates by sticking to RNA, we pre-incubated cell extracts with RNase A or with a 442 nucleotide fragment of the AC6 3’UTR that specifically binds to Bicc1 KH domains (Piazzon et al., 2012). Both RNase A treatment and the addition of synthetic target RNA each tended to increase HA-Bicc1 sedimentation, rather than inhibiting it (**Figure 1 - figure supplement 1E**). Likewise, when all KH domains in HA-Bicc1 were mutated specifically in their GXXG motifs (where at least one X must be a basic residue to allow RNA binding), it sedimented no less efficiently than wild-type (**Figure 1 - figure supplement 1F**). These data show that Bicc1 sedimentation in cell extracts is driven by one or several KH domains and independently of RNA binding, rather than by the SAM domain that drives Bicc1 polymerization in the cytoplasm.

**Figure 1.**
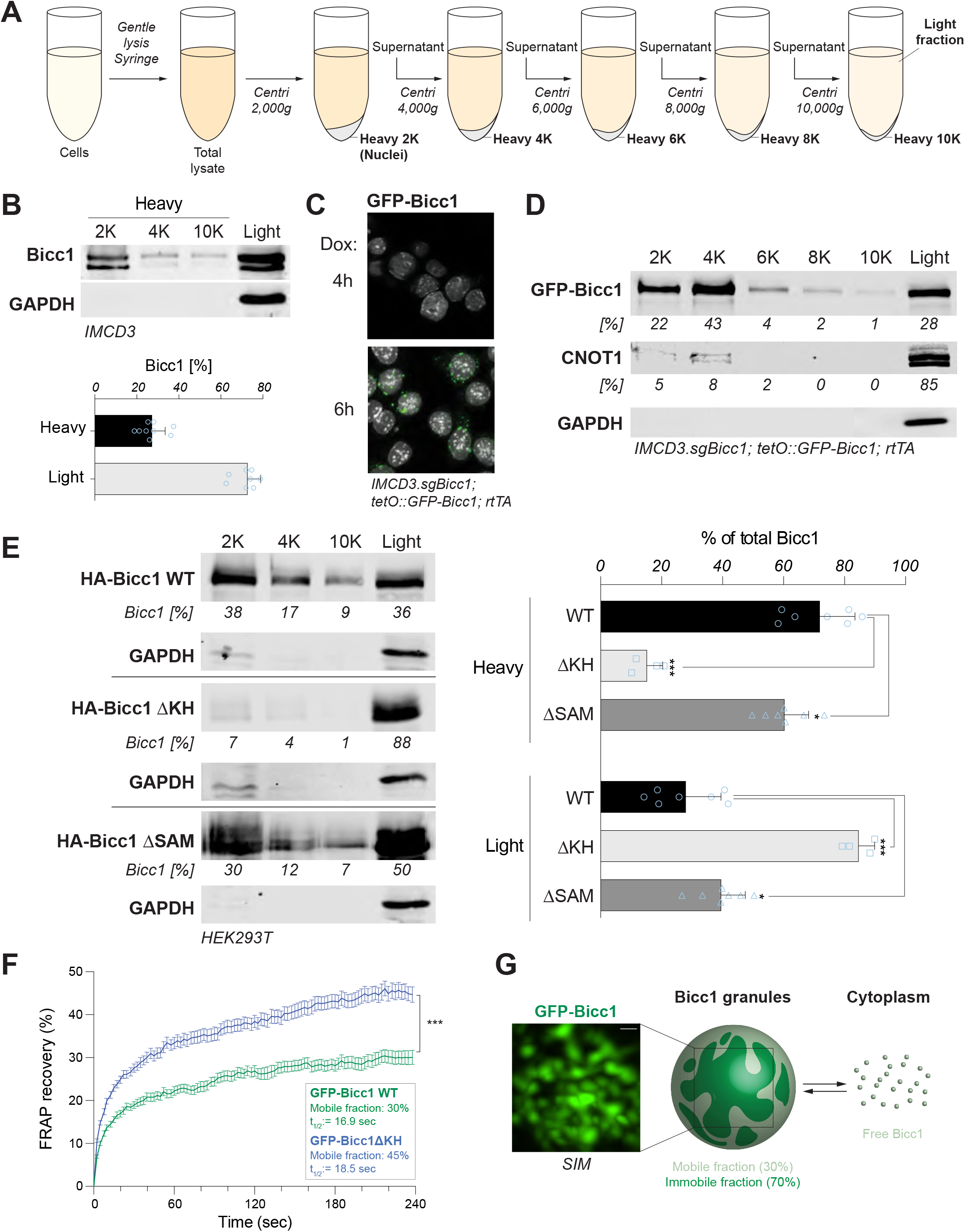
Sedimentation of Bicc1 by low-speed centrifugation is mediated by KH domains independently of their interaction with RNA. **(A)** Procedure of cell fractionation by low speed differential centrifugation. (**B**) Western blot of murine inner medullary collecting duct (IMCD3) cell extracts after stepwise centrifugation at 2,000 to 10,000 g as in (A). The relative distribution of mBicc1 in heavy (2K, 4K and 10K) versus light fractions is quantified in the graph on the right. Equal volumes of the indicated solubilized pellets and supernatant were analysed. Data are means + SD from nine independent experiments. **(C)** Imaging and (**D**) Western blots of a stepwise centrifugated cytoplasmic extract of an sgBicc1-mutated IMCD3 clone expressing doxycycline-inducible lentiviral *GFP-Bicc1*. Percentages below the gels in (D) are relative to the total. Dense fractions containing phase-separated protein granules are marked by the P-body component CNOT1. GAPDH marks the light fraction. **(E)** Sedimentation of HA-Bicc1 WT, ΔKH and ΔSAM by stepwise centrifugation of cell extracts of transiently transfected HEK293T cells. **(F)** Fluorescence-Recovery After Photobleaching (FRAP) of GFP-Bicc1 WT and ΔKH granules in HeLa cells. Results are mean values from 47 individual granules. Error bars show SEM. **(G)** A representative GFP-Bicc1 granule imaged by 3D SIM (left), and schematic representation of how its internal heterogeneity might be related to the protein dynamics observed in (F). Scale bar, 300 nm.

### The internal structure of the Bicc1 granules is heterogenous and exhibits gel-like features

To assess the contribution of KH domains to Bicc1 phase separation *in vivo*, we first imaged GFP-Bicc1 with or without KH domains by confocal microscopy in HeLa cells. In the most brightly fluorescent 70% of 125 cells examined, WT GFP-Bicc1 coalesced mostly in only a few large condensates, whereas in the remaining 30% marked by a 3-fold lower mean fluorescence intensity, GFP-Bicc1 granules remained small but were more numerous (**Figure 1 - figure supplement 2A**). By contrast, GFP-Bicc1 ΔKH failed to form large condensates and instead partitioned between small granules (42%) or fibrillar structures (22%) or a combination of both (34%). In good agreement, 3D reconstruction of structured illumination microscopy (SIM) images at high resolution confirmed that WT GFP-Bicc1 granules coalesce in space-filling interconnected meshworks, whereas GFP-Bicc1 ΔKH polymers extend longitudinally as fibrils (**Figure 1 - figure supplement 2B**). To address whether the KH repeat influences polymer dynamics, we measured the fluorescence recovery after photobleaching (FRAP) of GFP-Bicc1 WT and ΔKH in HeLa cells. Interestingly, the maximal fluorescence recovery was only 30% in GFP-Bicc1 WT compared to 45% in ΔKH, and it was reached quickly with half times less than 20 sec for both constructs (**Figure 1F**). Thus, a minority of Bicc1 within granules behaves as a liquid that rapidly exchanges molecules with the surrounding cytoplasm, whereas molecular diffusion of the larger immobile fraction is limited by a more rigid, gel-like structure, in part due to the KH repeat (**Figure 1G**). Taken together, these results demonstrate that the KH repeat modulates both the 3D configuration and the dynamics of Bicc1 granules.

### Both the SAM and KH domains self-interact and contribute to cytosolic Bicc1 condensation

Bicc1 can self-associate with both its SAM domain and with the KH tandem repeat in yeast two-hybrid experiments, but no interactions have been observed between KH domains (Rothé et al., 2018). To evaluate KH domain self-interactions by an alternative approach, we loaded glutathione sepharose beads with GST fusion proteins of the entire KH repeat or of individual KH domains and incubated them with HA-Bicc1 in HEK293T cell extracts. Western blot analysis of bound proteins revealed that GST-KH and GST-KH_1_ pulled down full-length and ΔSAM mutant HA-Bicc1, but only minimal amounts of HA-Bicc1 ΔKH (**Figure 2 - figure supplement 1A**). Participation of KH_1_ in an intermolecular KH:KH interface is consistent with the X-ray structure of isolated BICC1 KH_1_ where an extended β-sheet stabilized by contacts between β1 strands mediates homo-dimerization (PDB: 6GY4) (**Figure 2 - figure supplement 1B**). Superimposition of this human KH_1_ domain structure on a molecular model of the KH tandem repeat indicates that the KH_1_-KH_1_ interface combines all RNA binding clefts as an extended platform on a single surface accessible to RNA (**Figure 2 - figure supplement 1C**). In contrast to what we observed with KH_1_, pull-down of the KH repeat by KH_2_ was close to background seen with GST alone, and the KH_3_ and the KH-like 1 or 2 domains individually or in combinations remained insoluble. HA-Bicc1 and truncation constructs lacking KH or SAM domains also bound only weakly to a fusion of GST with the Bicc1 IVS. These results suggest that Bicc1 polymers are jointly organized by SAM:SAM association and by KH_1_ binding to the KH tandem repeat.

To test whether the KH repeat cooperates with SAM domain polymerization *in vivo*, we analyzed the relative contributions of each domain to the co-assembly of HA-Bicc1 granules with cytoplasmic GFP-Bicc1. While GFP-Bicc1 co-immunoprecipitated with HA-Bicc1 in HEK293T cell extracts as expected, disruption of the SAM:SAM interface by alanine substitutions in residues D913, K915 and E916 of HA-Bicc1 (mutation D) (Rothé et al., 2015) or deletion of the KH tandem repeat each reduced this interaction by 60-75% (**Figure 2A**). HA-Bicc1 and GFP-Bicc1 granules also co-localized in the cytoplasm of transfected HeLa cells (**Figure 2B**). Importantly, despite their weakened affinity for GFP-Bicc1, both HA-Bicc1ΔKH and the polymerization-deficient HA-Bicc1 mutD were still recruited to cytoplasmic GFP-Bicc1 condensates. By contrast, HA-Bicc1ΔKH-mutD lacking both the KH tandem repeat and the SAM:SAM polymerization interface failed to bind to and co-localize with GFP-Bicc1 (**Figure 2A-B**). These data agree with our pull-down assays that the KH repeat increases the valency of Bicc1 self-interactions in cytoplasmic granules.

**Figure 2.**
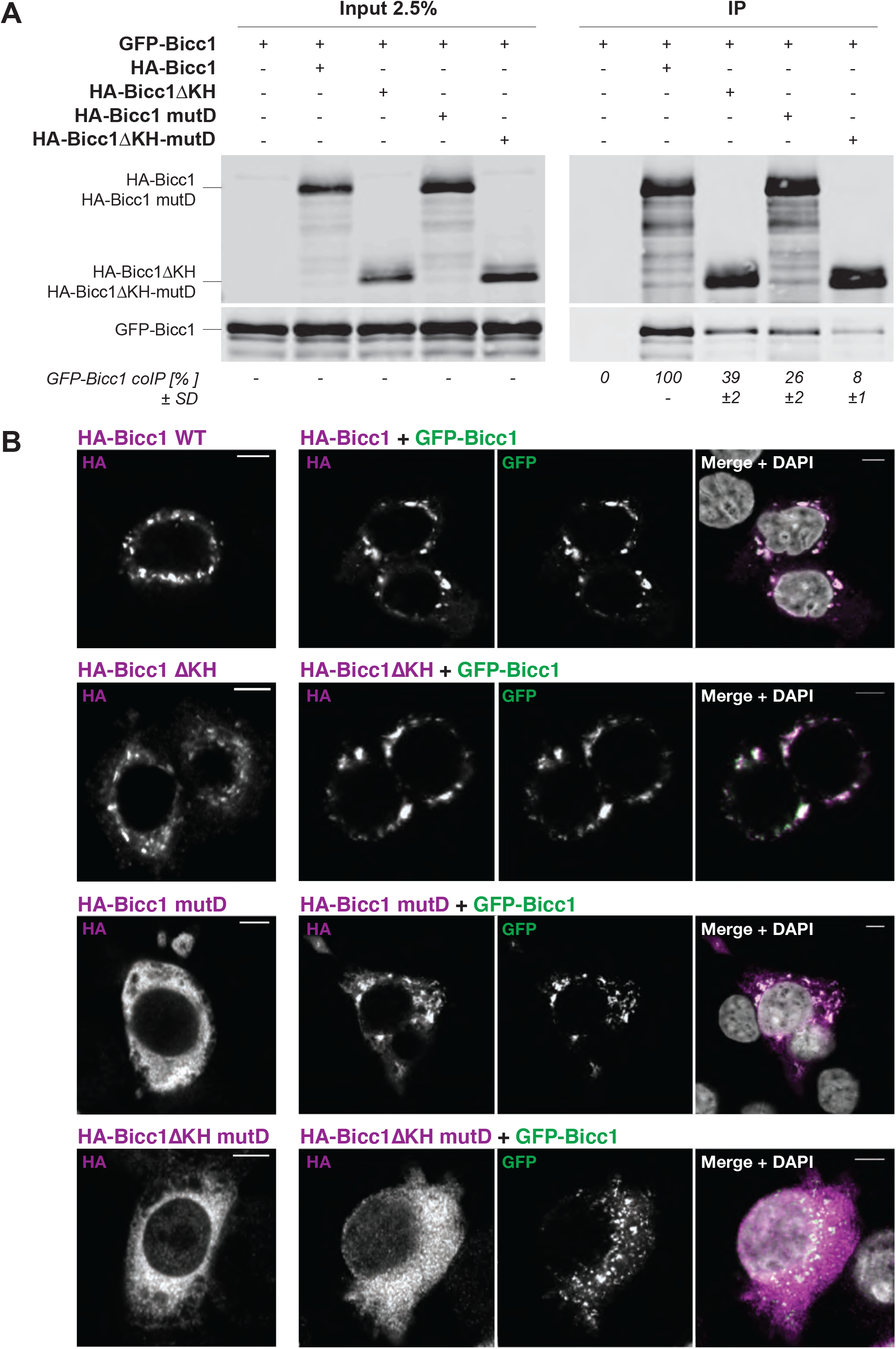
KH domain self-interactions cooperate with SAM polymerization to condensate Bicc1 in cytoplasmic granules. **(A)** Co-immunoprecipitation of GFP-Bicc1 with HA-Bicc1 WT, ΔKH, mutD or ΔKH-mutD co-expressed in HEK293T cells. The amount of co-immunoprecipitated GFP-Bicc1 was normalized over the input and expressed relative to the WT HA-Bicc1 condition. Values below the gel are means ± SD of two independent experiment. They represent the ratio of co-immunoprecipitated GFP-Bicc1 to the input and are expressed relative to the condition with WT HA-Bicc1. **(B)** Immunofluorescence staining of GFP-Bicc1 and HA-Bicc1 WT, ΔKH, mutD or ΔKH-mutD co-expressed in HeLa cells. Scale bars, 5 µm.

### Bicc1 granules promote the phase separation of target mRNAs

To test whether Bicc1 self-interactions may mediate phase separation of bound mRNAs, we imaged GFP-Bicc1 and its association with 12xMS2-tagged reporter mRNAs in real time by co-expressing a Halo-tagged tandem repeat of MS2 coat protein fused to a nuclear localization sequence (2xMCP-Halo-NLS). In this setting, the localization of MS2-tagged mRNA in the cytoplasm can be visualized by labelling bound 2xMCP-Halo-NLS with the fluorescent Halo-tag ligand TMR (Halstead et al., 2015) (**Figure 3A**). Irrespective of the presence or absence of GFP-Bicc1, empty-12xMS2 control mRNA lacking a Bicc1 binding site diffused throughout the cytoplasm as expected (**Figure 3B**). By contrast, Dand5-3’UTR-12xMS2 reporter containing the Bicc1-specific binding motif GACGUGAC of the 3’UTR of Dand5 mRNA (Minegishi et al., 2021) only diffused freely in cells without Bicc1, whereas coxpression of GFP-Bicc1 led to its concentration in cytoplasmic condensates (**Figure 3C**). To test whether Bicc1 binding forms similar condensates of its own mRNA (Chicoine et al., 2007; Piazzon et al., 2012) and to study this process in real time, we inserted the 12xMS2-tag in the GFP-Bicc1 construct. Thus, during live imaging, all cells expressing GFP-Bicc1 protein also transcribe the target mRNA. TMR labelling of 2xMCP-Halo-NLS revealed that the 12xMS2-tagged GFP-Bicc1 mRNA accumulated in GFP-Bicc1 granules (**Figure 3D**). Furthermore, this co-localization persisted in the presence of puromycin (**Figure 3E**), indicating that it is independent of ongoing translation. To assess the dynamics of the RNA fraction in Bicc1 granules, we performed FRAP analysis of the TMR signal. We observed 50% recovery within 4 min (**Figure 3F**), consistent with a phase transition of the mRNA alongside GFP-Bicc1 protein condensates. Taken together, these data suggest that Bicc1 partly immobilizes associated transcripts in gel-like condensates and partly exchanges them for mRNA in the surrounding fluid phase.

**Figure 3.**
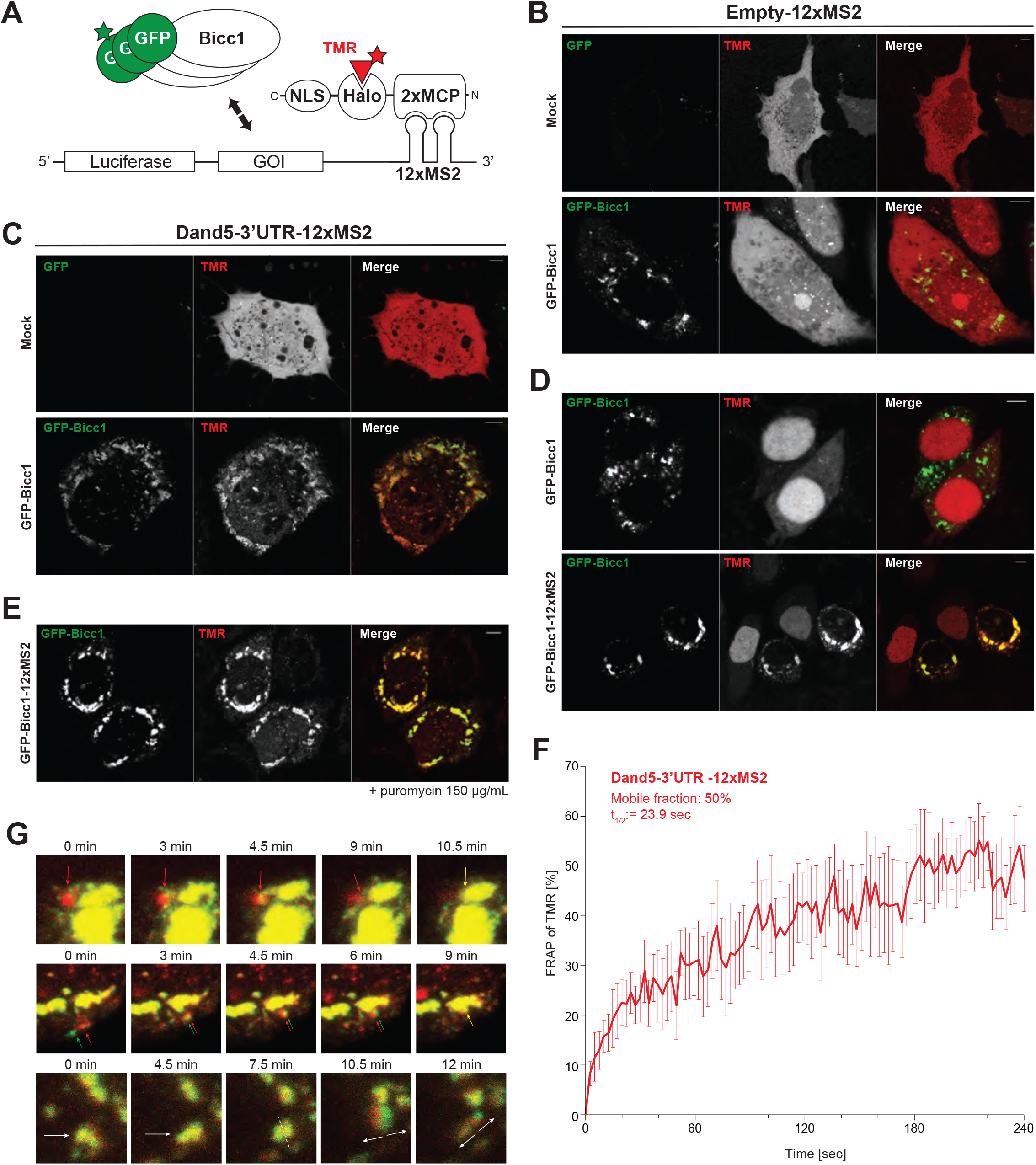
Bicc1 promotes the phase separation of target mRNAs in cytoplasmic granules. **(A)** Methodology to image the distribution of 12xMS2-tagged luciferase reporter mRNAs and its regulation by GFP-Bicc1. Halo-tagged MS2-coat protein (MCP) fused to a Nuclear Localization Signal (NLS) normally accumulates in the nucleus, but when bound to a tandem repeat of 12 MS2 hairpins in reporter mRNAs of genes of interest (GOI), such as luciferase fused to the Bicc1 coding sequence (CDS) or to a Dand5 3-’UTR fragment, MCP fusion protein is retained in the cytoplasm. HaloTag is labeled using the fluorescent ligand TMR. Luciferase alone (Empty-12xMS2) is a negative control. **(B-C)** Live imaging of GFP-Bicc1 fluorescence, together with HaloTag/TMR-labelled Empty-12xMS2 (B) or Dand5 3’UTR-12xMS2 reporter mRNAs (C). **(D-E)** Live imaging of GFP-Bicc1 (green) and its own 12xMS-tagged mRNA (red) before (D) and after treatment with puromycin for 30 min (E). **(F)** Fluorescence-Recovery After Photobleaching (FRAP) of the Dand5-3’UTR reporter mRNA present in GFP-Bicc1 granules of HeLa cells and marked by TMR bound to 2xMCP-Halo protein. Results are mean values from 8 individual granules. Error bars show SEM. **(G)** Time-lapse imaging of fusion (top and mid lanes) and scission (bottom lane) events between mRNA foci and GFP-Bicc1 granules.

### The Bicc1 granules can acquire actomyosin-dependent mobility

To monitor the dynamics of mRNA translocation to cytoplasmic Bicc1 granules in real time, we tracked GFP and TMR fluorescence in time-lapse movies. Already at the onset of GFP fluorescence, the largest Bicc1 condensates contained copious amounts of mRNA and were poorly mobile but continued to irreversibly fuse with more mobile Bicc1-free mRNA particles passing in their vicinity (**Figure 3G, Figure 3 - video 1-3**). Alternatively, during spikes of directional movement, GFP-Bicc1 protein granules were observed to split (**Figure 3 - figure supplement 1A-C**) or, in rare instances, to dissociate from mRNA particles (**Figure 3G, Figure 3 – figure supplement 1D, Figure 3 - video 4-5**). To test whether this mobility depends on the cytoskeleton, we tracked the trajectories of GFP-Bicc1 granules in cells treated with the Myosin II inhibitor Blebbistatin or with the ROCK inhibitor Y27632. In the presence of either of these drugs, the total distance travelled by individual GFP-Bicc1 granules decreased by 40-50% (**Figure 3 - figure supplement 1E**). The mobility of GFP-Bicc1 granules also decreased on average by 12.5 ± 2.2% upon treatment with the actin polymerization inhibitor cytochalasin D, and by 15.3 ± 2.8% after microtubule-depolymerisation by Nocodazole. By contrast, treatment with the kinesin inhibitor Monastrol had no effect. These observations suggest that Bicc1 condensates are eligible to cytoskeletal transport mediated by actomyosin and to a lesser extent by microtubules.

### ANKS3 binds the KH domains of Bicc1 and competes with RNA binding

The Bicc1 SAM domain and the KH repeat can also each bind to ANKS3 (Rothé et al., 2018). To map where ANKS3 contacts the KH tandem repeat, we compared binding to individual Bicc1 domains in yeast two-hybrid (Y2H) assays, including KH_1_, KH_2_ or a combination of KHL_1_- KH_3_-KHL_2_ domains (Bicc1-KHL). The strength of interaction was assessed by titrating 3- aminotriazol (3AT), a competitive inhibitor of the reporter gene product. Binding of ANKS3 to the KH tandem repeat resisted up to 60 mM 3AT, whereas among KH fragments, only Bicc1-KHL bound ANKS3 above background levels, and this interaction was lost already at 0.5 mM 3AT (**Figure 4A**). Furthermore, ANKS3 binding to the Bicc1 SAM domain or to itself (Gal4-AD-ANKS3) was inhibited by as little as 5-10 mM 3AT. To independently validate the contribution of individual KH domains, we incubated ANKS3-Flag in cell extracts with the entire KH tandem repeat, or only the KH_1_ or KH_2_ domains, or with the IVS fused to GST on glutathione sepharose beads. Western-blot analysis of bound proteins revealed that only the complete KH tandem repeat retained ANKS3-Flag (**Figure 4B**). These results suggest that tight binding of Bicc1 to ANKS3 requires an extended surface of more than one KH domain.

**Figure 4.**
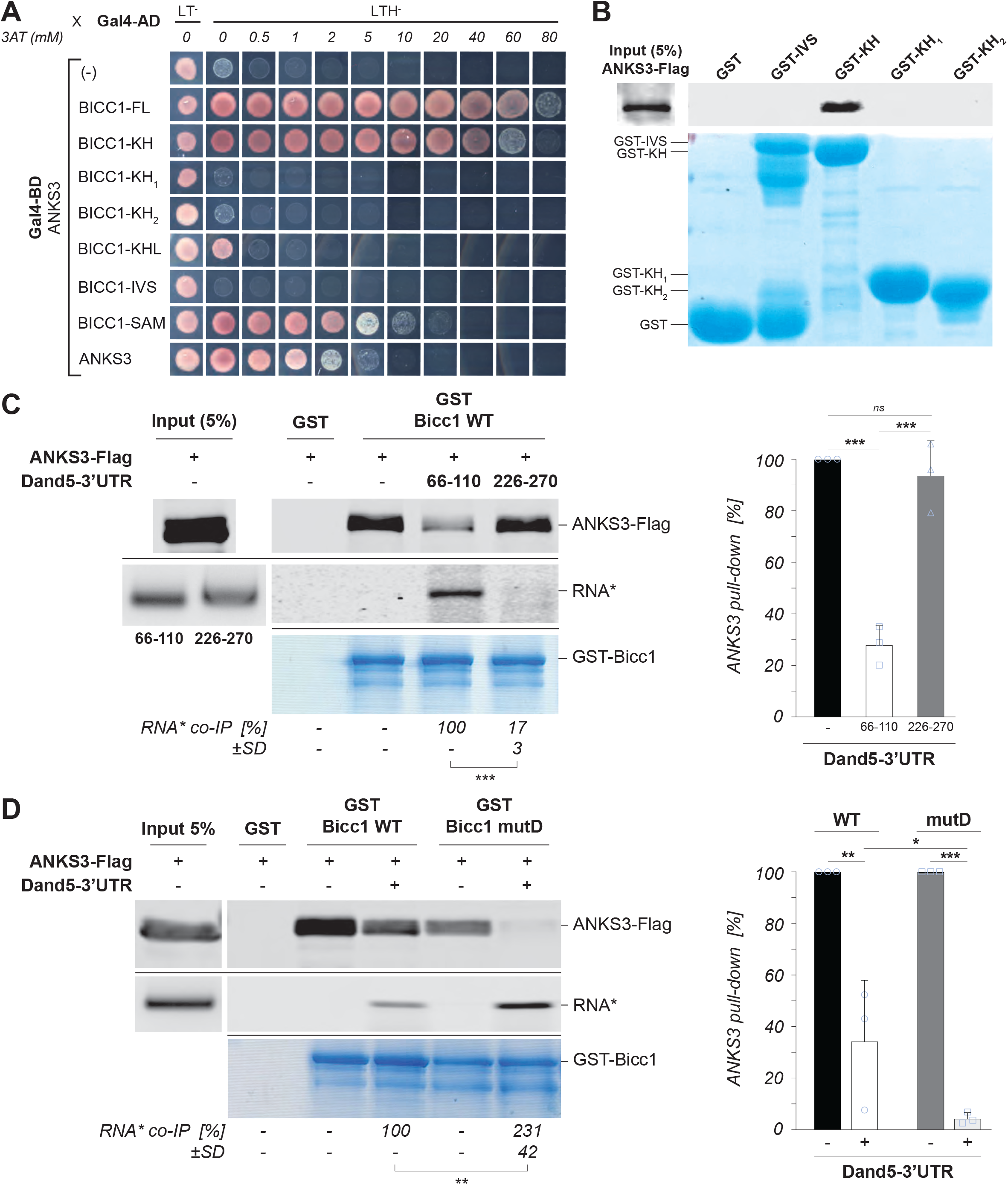
ANKS3 binds to the Bicc1 KH domains and competes with RNA binding. **(A)** Yeast two-hybrid mapping of the interaction between a fusion of ANKS3 with the DNA-binding domain of Gal4 (Gal4-BD) and human BICC1 KH domains fused to Gal4 activation domain (Gal4-AD). Controls in non-selective medium without leucine (L) and tryptophan (T) are shown in the first column (LT^−^). Interactions were revealed at the indicated concentrations of 3-aminotriazole (3AT) in triple selective medium (LTH^−^) lacking histidine. **(B)** Pull-down of ANKS3-Flag from HEK293T cell extracts by glutathione sepharose beads coated with recombinant domains of Bicc1 fused to GST. **(C)** ANKS3 and specific target RNA compete for Bicc1 binding. Western blot analysis of ANKS3-Flag from HEK293T cell extracts before (input) and after GST pull-down by ribonucleoparticles (RNP) of full-length Bicc1 (GST-Bicc1) that were pre-loaded or not with saturating amounts of the fluorescently labelled Dand5-3’UTR RNA fragment 66-110 or, as a control for non-specific binding, fragment 226-270 (*). The fluorescence of the RNA (middle panels), and Coomassie blue staining of GST-KH protein retained by the beads (bottom panel) are shown below. **(D)** Western blot analysis of ANKS3-Flag from HEK293T cell extracts before (input) and after GST pull-down by *in vitro* reconstituted ribonucleoparticles (RNP) of recombinant GST-Bicc1 full-length WT or mutD that were pre-loaded or not with saturating amounts of the fluorescently labelled Dand5-3’UTR RNA fragment 66-110. The fluorescence of the RNA (middle panels), and Coomassie blue staining of GST-KH protein retained by the beads (bottom panel) are shown below. In (C-D), the amounts of target RNA (*) and of ANKS3-Flag retained by the beads are shown below the gels and in the histogram to the right, respectively. Values are relative to the pull-down with wild-type (WT) Bicc1 (100%). Data are means + SD from three independent experiments. ns: non-significant, *p < 0.05, **p < 0.01, ***p < 0.001 (Student’s t-test).

To assess whether ANKS3 and RNA compete for Bicc1 binding, we performed GST pull-down in the presence of a Dand5-3’UTR_66-110_ RNA fragment that binds KH_1_ and KH_2_ domains with high affinity, or with a similarly sized non-specific Dand5-3’UTR_226-270_ fragment that lacks Bicc1 binding sites (Minegishi et al., 2021). We found that the saturation with the 66-110 transcript inhibited the pull-down of ANKS3-Flag by GST-Bicc1 by 70%, whereas no significant inhibition was observed with the non-specific 226-270 fragment (**Figure 4C**). Interestingly, saturation with Dand5-3’UTR_66-110_ further reduced the pull-down of ANKS3-Flag by polymerization-deficient mutant Bicc1 mutD to nearly background levels (**Figure 4D**). Conversely, the retention of RNA by mutD versus WT Bicc1 increased more than twofold. These data show that binding of Bicc1 to ANKS3 or RNA is mutually exclusive, and that SAM:SAM interactions bias this competition in favor of ANKS3.

### Endogenous ANKS3 buffers the binding of Bicc1 to specific target mRNAs in IMCD3 cells

To test whether ANKS3 and RNA compete for Bicc1 binding *in vivo*, we analyzed the recruitment of endogenous *Bicc1* and *Adcy6* mRNAs to Bicc1 RNPs in IMCD3 cells expressing doxycycline-inducible ANKS3 shRNA (Schlimpert et al., 2018). Validation by reverse transcription (RT) and quantitative polymerase chain reaction (qPCR) analysis confirmed that doxycycline administration depleted ANKS3 mRNA by >80% (**Figure 5 - figure supplement 1A**), and without changing Bicc1 protein levels (**Figure 5 - figure supplement 1B**), or the expression of β-actin control mRNA, or the background of its non-specific binding in anti-Bicc1 immunoprecipitates (**Figure 5 - figure supplement 1C**). However, co-IP of *Adcy6* and *Bicc1* transcripts with Bicc1 protein increased on average 5-fold in ANKS3-depleted cells compared to control. Unexpectedly, Bicc1 mRNA also increased 7-fold in the input fraction, whereas Adcy6 mRNA expression levels were unchanged. We conclude that ANKS3 depletion selectively stabilizes Bicc1 binding to *Adcy6* mRNA, whereas increased binding to *Bicc1* transcripts is accompanied by a proportionate net increase of Bicc1 mRNA.

To test whether ANKS3 similarly inhibits Bicc1 binding to mRNA in gain-of-function experiments, we co-transfected HA-Bicc1 and dsVenus-*Dand5*-3’UTR reporter with ANKS3-Flag or empty vector control in HEK293T cells (Minegishi et al., 2021). RT-qPCR analysis of anti-HA immunoprecipitates confirmed that HA-Bicc1 specifically enriches the dsVenus-*Dand5*-3’UTR mRNA on average more than 25-fold relative to the background, whereas no enrichment was observed for the β-actin control mRNA (**Figure 5A, Figure 5 - figure supplement 1D**). Furthermore, co-expression with ANKS3-Flag reduced this specific association of HA-Bicc1 with target mRNA by about 80% (**Figure 5A**). Co-immunoprecipitation of HA-Bicc1 transcripts decreased to a similar extent, confirming that ANKS3 inhibits RNA binding.

**Figure 5.**
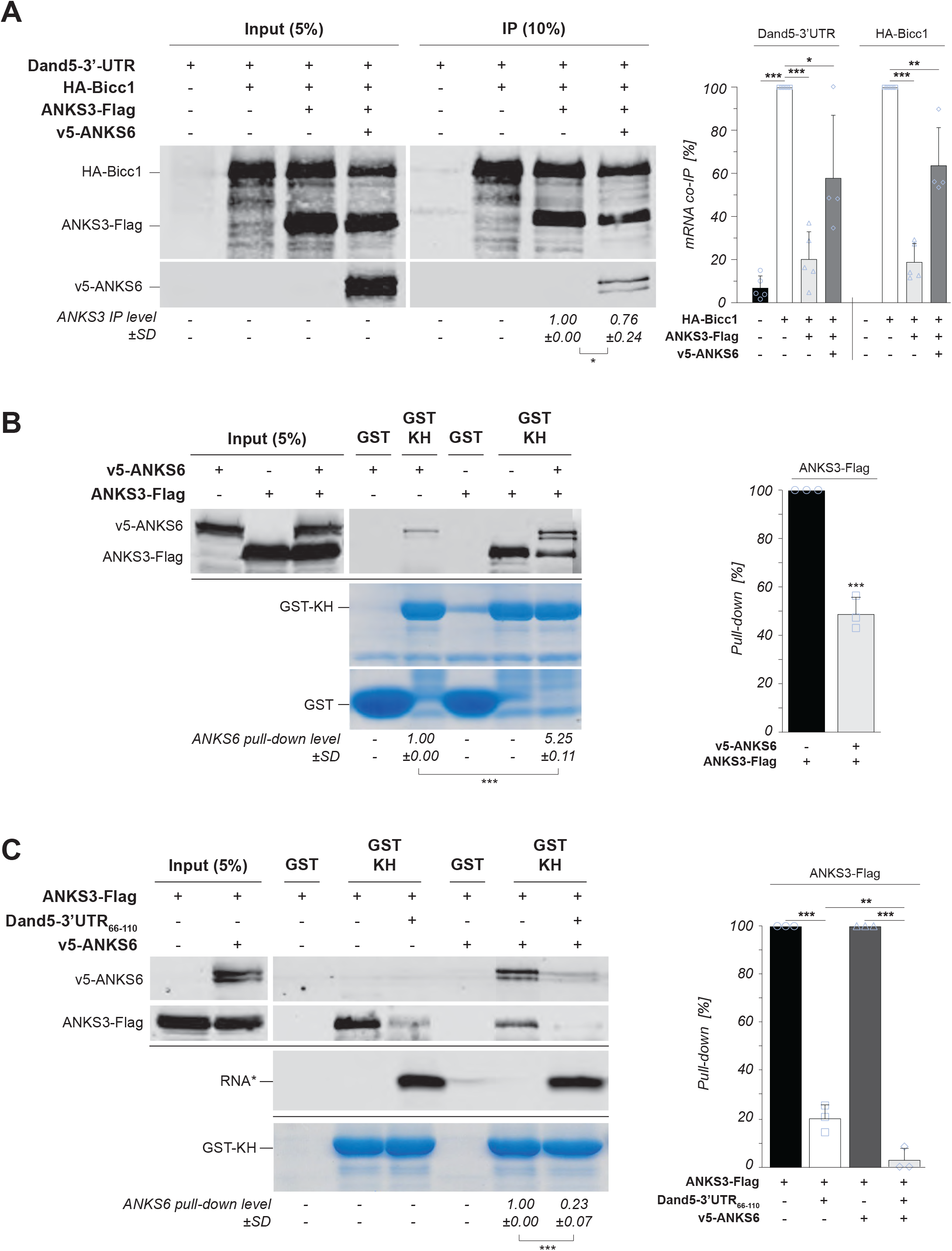
ANKS6 augments Bicc1 binding to target RNAs while destabilizing the ANKS3-KH interaction. **(A)** RNA co-immunoprecipitation with HA-Bicc1 from cytoplasmic extracts of HEK293T cells expressing the dsVenus-Dand5-3’UTR reporter and HA-Bicc1 alone or in combination with ANKS3-Flag and v5-ANKS6. Western blots of protein inputs and the level of ANKS3 in the IP fraction relative to immunoprecipitated Bicc1 are shown on the left. On the right, RT-qPCR analysis shows the ratios of co-immunoprecipitated Dand5-3’UTR reporter mRNA and HA-Bicc1 transcript normalized to their amounts in inputs and to HA-Bicc1 protein in the IP, relative to the corresponding condition with HA-Bicc1 only. **(B)** Left: GST pull-down of ANKS3-Flag in HEK293T cell extracts by GST-Bicc1-KH in presence or absence of v5-ANKS6. Right: Quantification of bound ANKS3-Flag relative to the pull-down without ANKS6 (100%). **(C)** GST pull-down as in (B), but using GST-Bicc1-KH beads that were first saturated with fluorescently labelled synthetic Dand5-3’UTR_66-110_ RNA where indicated. Data are means + SD from at least three independent experiments. ns: non-significant, *p < 0.05, **p < 0.01, ***p < 0.001 (Student’s t-test).

### Sequestration of ANKS3 by ANKS6 stimulates Bicc1 binding to mRNA

To investigate how Bicc1 KH domains are freed from ANKS3 inhibition to bind mRNA, we tested whether ANKS3 is regulated by ANKS6. Interestingly, co-expression of v5-ANKS6 depleted ANKS3-Flag in Bicc1 immunoprecipitates by nearly 25% and restored significant HA-Bicc1 binding to both the *Dand5* reporter mRNA and to HA-Bicc1 transcripts (**Figure 5A**). To validate that ANKS6 neutralizes ANKS3, we also tested its effect on ANKS3-Flag and mRNA binding to GST-KH. We found that even though the presence of v5-ANKS6 in cell extracts reduced the GST-KH pull-down of ANKS3-Flag by half, the residual bound ANKS3-Flag in turn increased the retention of V5- ANKS6 on GST-KH beads (**Figure 5B**), consistent with its role in recruiting ANKS6 to Bicc1 (Rothé et al., 2018). To assess the consequences for RNA binding, we performed a competition assay by saturating the GST-KH beads with the fluorescent Dand5-3’UTR_66-110_ RNA. While saturation of GST-KH with target RNA diminished the pull-down of ANKS3-Flag by 80 ± 6%, the presence of v5- ANKS6 further reduced it close to background levels (**Figure 5C**). These data agree with the conclusion of our coimmunoprecipitation assays that recruitment of ANKS6 frees Bicc1 KH domains for increased mRNA binding by weakening their interaction with ANKS3.

### ANKS3 and ANKS6 spatiotemporally couple RNA binding to Bicc1 phase transitioning

When expressed without ANKS proteins, Bicc1 can bind RNA independently of its SAM domain (Piazzon et al., 2012). Therefore, and since emulsification by ANKS3 or its inhibition by ANKS6 inhibited or stimulated RNA binding, respectively, we wondered whether ANKS proteins spatiotemporally couple RNA binding of Bicc1 to its phase separation. To address this, we imaged the recruitment of GFP-Bicc1-12xMS2 mRNA to GFP-Bicc1 granules in HeLa cells expressing ANKS3-Flag alone or together with V5-ANKS6. In live imaging, only GFP-Bicc1 and its TMR-labelled mRNA are visible. Therefore, as a control, we validated in fixed cells that co-expression with ANKS3-Flag alone dispersed large GFP-Bicc1 condensates (**Figure 6 - figure supplement 1A-B**), whereas co-expression with ANKS3-Flag plus V5-ANKS6 blocked this effect and rescued the assembly of all three proteins in large condensates as expected (Rothé et al., 2018) (**Figure 6 - figure supplement 1C**). Since GFP-Bicc1 was too widely dispersed in ANKS3-expressing cells to estimate its colocalization with mRNA in entire cells, we instead tracked discrete RNPs over time. In the absence of ANKS3-Flag, large GFP-Bicc1 granules remained stably associated with reporter mRNA (**Figure 6A, Figure 6 - video 1**), whereas in its presence, the peaks of GFP and TMR fluorescence strongly anti-correlated during at least 10 min (**Figure 6B, Figure 6 - video 2**). Thus, ANKS3 potently unmixes GFP-Bicc1 and mRNA foci. Conversely, the rescue of large GFP- Bicc1 condensates by ANKS6 restored their stable co-localization with GFP-Bicc1 mRNA (**Figure 6C, Figure 6 - video 3**). Importantly, co-recruitment of V5-ANKS6 also rescued large condensates of ANKS3-Flag with HA-Bicc1 ΔKH that cannot bind RNA (**Figure 6 - figure supplement 1C**), confirming that increased RNA binding is not the cause of ANKS6-mediated granule formation, but rather its consequence. Overall, these data indicate that binding of Bicc1- ANKS3 complexes to specific mRNAs is coupled to the rescue of their phase transitioning by ANKS6 (**Figure 7**).

**Figure 6.**
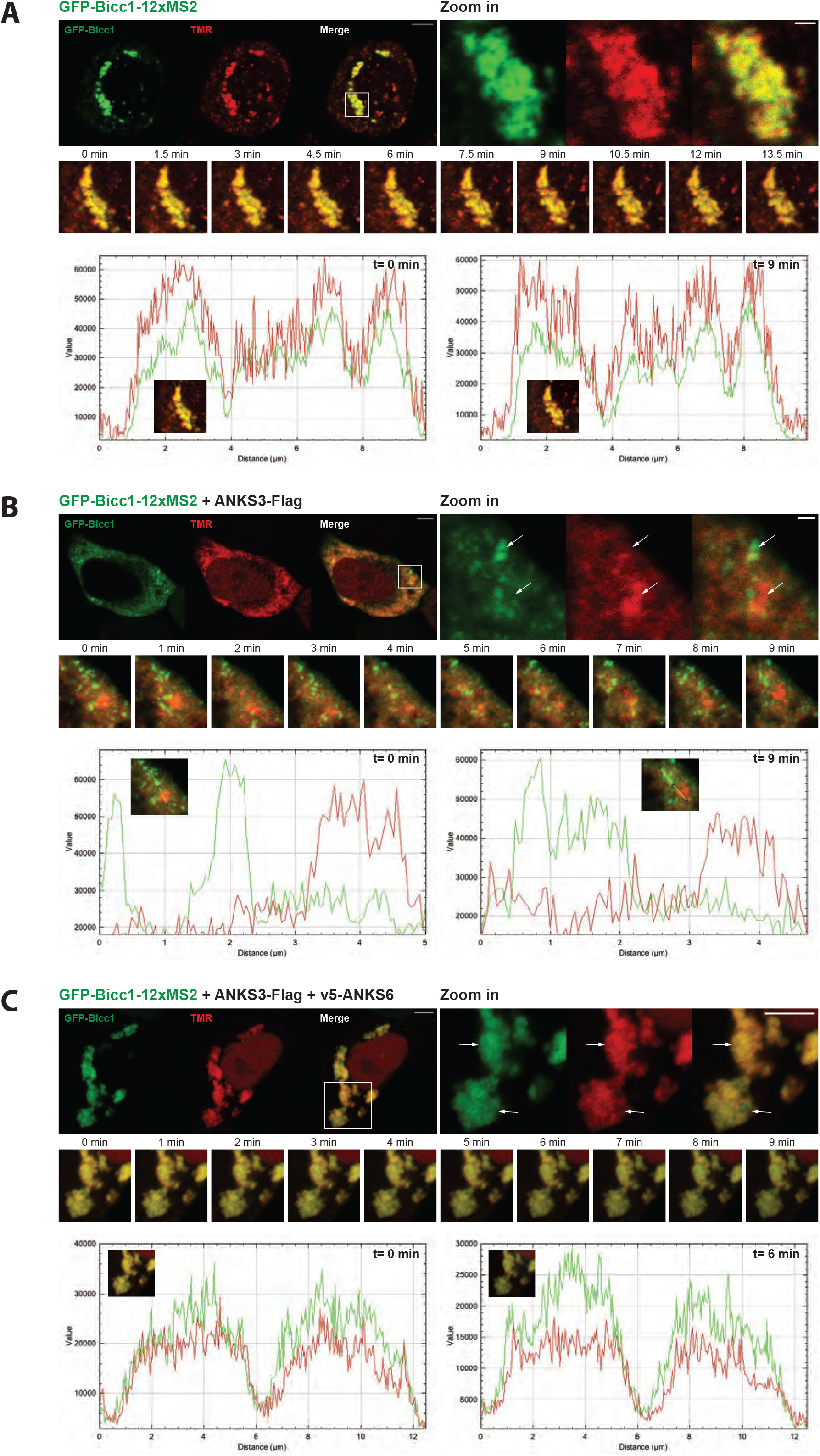
Bicc1 phase separation and RNA binding are connected through ANKS3 and ANKS6. **(A-C)** Live imaging of GFP-Bicc1 (green) and its own 12xMS2-tagged mRNA (red) in HeLa cells as in figure 4. GFP-Bicc1-12xMS2 was expressed alone (A), or in combination with ANKS3- Flag (B), or ANKS3-Flag and v5-ANKS6 (C). The fluorescent signal plots were generated using the multichannel plot profile analysis in Image J.

**Figure 7.**
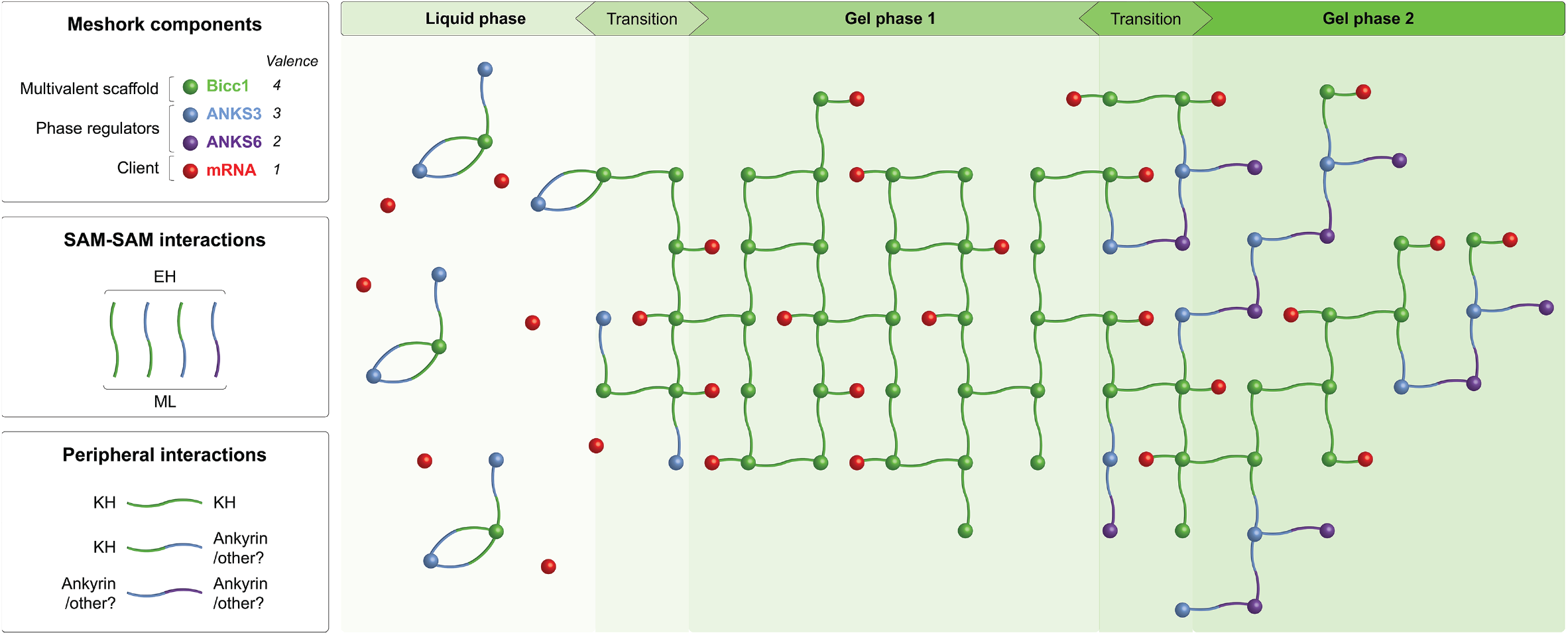
Schematic of the influence of phase-transitioning on the mRNA binding activity of Bicc1, and its dual regulation by ANKS3 and ANKS6. Topological remodelling of multivalent protein-protein interactions in the Bicc1:ANKS3:ANKS6 meshwork favors the transition between liquid (Bicc1:ANKS3) and gel-like metastable states (RNA:Bicc1, or RNA:Bicc1:ANKS3:ANKS6) to either prevent (left) or promote specific binding of Bicc1 to mRNA (right), depending on the relative concentrations of individual components (spheres) and the number of their binding sites (valence) that engage in homo- or heterotypic interactions (boxes). Vertical connectors represent homo- or heterotypic SAM:SAM interactions horizontal connectors represent interactions by KH domains, Ankyrin repeats and additional interacting protein domains.

## DISCUSSION

While ANKS3 can disperse Bicc1 self-polymers, co-recruitment of ANKS6 by ANKS3 has recently been shown to rescue their condensation in large cytoplasmic granules (Rothé et al., 2018). Prompted by these observations, we here asked whether phase transitioning and its regulation by ANKS3 and ANKS6 influence Bicc1 binding to specific target mRNAs or *vice versa*. We found that self-polymerization by the SAM domain nucleates phase transitioning of Bicc1 and associated transcripts into cytoplasmic RNPs that can acquire cytoskeleton-mediated mobility and split or fuse with each other. Immunofluorescent staining and co-immunoprecipitation analysis of reconstituted Bicc1 oligomers revealed that their clustering in cytoplasmic condensates additionally involves KH domains independently of their RNA-binding GXXG motifs. Moreover, yeast-two- hybrid and RNA co-immunoprecipitation assays and pull-down of *in vitro* reconstituted RNPs revealed that the KH tandem repeat also directly interacts with ANKS3, and that ANKS3 in turn reduces Bicc1 binding to specific target mRNAs. By contrast, ANKS6 facilitated mRNA binding by destabilizing the interaction of ANKS3 with the KH domains. Overall, these observations suggest that the multivalent interactions with ANKS3 and the co-recruitment of ANKS6 synchronize and spatially couple specific RNA binding of Bicc1 with its phase separation.

### KH domain self-interactions and SAM domain polymerization jointly shape higher order Bicc1 condensates

A significant fraction of endogenous Bicc1 in IMCD3 cell extracts sedimented at low centrifugation speed. Our analysis of Bicc1 truncation mutants revealed that *in vitro* sedimentation mainly required the KH tandem repeat, but not its RNA-binding GXXG motifs. By contrast, Bicc1 condensation in the cytoplasm is nucleated primarily by the SAM domain (Maisonneuve et al., 2009; Rothé et al., 2015), suggesting that the relative contributions of KH and SAM domains to promote granule formation are context-dependent. Indeed, increasing the salt concentration restored Bicc1 solubility during cell extraction. A similar effect of the ionic strength on phase separation has been described for other proteins (Qamar et al., 2018). In keeping with a role in stimulating Bicc1 condensation *in vivo*, we found that the KH repeat mediates the recruitment of truncated Bicc1 ΔSAM molecules into cytoplasmic GFP-Bicc1 co-polymers. Furthermore, our SIM images show that GFP-Bicc1 ΔKH polymers fail to coalesce in space-filling interconnected meshworks and instead form fibrillar structures. Together, these observations suggest that the Bicc1 SAM domain nucleates longitudinally growing fibrils that are remodeled through interconnecting KH domain self-interactions, presumably to stiffen intramolecular dynamics and thereby promote gel-like phase transitions. In line with this interpretation, our FRAP analysis revealed that 70% of GFP-Bicc1 within granules was immobile. Furthermore, deletion of the KH domains increased the mobile fraction from 30 to 45% at the expense of the static pool. The rapid exchange of the mobile fraction within seconds (t_1/2_ < 20 sec) resembles the dynamics of SGs and PBs that form by liquid-liquid phase transitioning (Aizer et al., 2014; Kedersha et al., 2000, 2005). However, the RNA-independent role of KH domains in shaping Bicc1 condensates sharply contrasts the phase transitioning of SG and PB components mediated by intrinsically disordered regions and RNA (Boeynaems et al., 2018). Remodeling of Bicc1 fibers by the structured KH domains into a more densely interconnected meshwork thus likely promotes optimal immobilization in gel-like granules.

### Bicc1 granule dynamics

Our live imaging experiments show that GFP-Bicc1 granules are mobile, with a subset showing directional movements driven by actomyosin and to a lesser extent by microtubules. In line with this observation, previous analysis of the Bicc1 protein interactome identified components of the actomyosin network, including Filamin, Alpha-actinin-4, Actin-like protein 6A and several ARHGEF and ARHGAP proteins (Leal-Esteban et al., 2018). By contrast, PBs use microtubules for long-range movements and remain stationary when associated with actin (Koppers et al., 2020). Spikes of directional motion of small GFP-Bicc1 granules and associated RNA coincided with fusion and fission, or with their incorporation into larger stationary Bicc1 RNPs. Immobilization likely involves crosslinking by KH domains, given the FRAP and SIM data discussed above. In good agreement, several other KH tandem repeats have been shown to crystallize as dimers or tetramers by helix-helix packing or β-sheet augmentation (Valverde et al., 2008) (**Figure 2 - figure supplement 1B**). Furthermore, our pull-down assays show that Bicc1 self-interactions involve at least the KH_1_ domain. Analogous experiments with the other KH and KH-like domains were inconclusive because their individual fusion to GST and deletions in full-length Bicc1 resulted in insolubility (unpublished observation). However, our molecular modeling predicts that self-dimerization rigidifies the Bicc1 KH_1_ in an extended RNA-binding platform (**Figure 2 - figure supplement 1C**), in sharp contrast to the apparent flexibility of KH_1_ in the monomeric structure of the Bicc1-related *C. elegans* protein GLD-3 (Nakel et al., 2010, 2015). Thus, rigidification of KH domains in interconnected protein meshworks likely influences their dynamics of RNA binding. For example, KH_1_ flexibility could be critical to help KH_2_ recognize specific target RNAs by prying open nearby secondary structures (Minegishi et al., 2021). Conversely, the arrangement in a rigid multimeric platform might stabilize the conformation of bound RNA.

### Regulation of Bicc1 phase separation and RNA binding by ANKS3 and ANKS6

While RNA frequently drives phase transitions of protein condensates, we found that neither RNA nor the RNA-binding GXXG motifs are necessary for the sedimentation *in vitro* or for granule formation of Bicc1 *in vivo*, respectively. Thus, the association of target mRNAs with Bicc1 granules is a consequence rather than a cause of their phase transitioning. To corroborate this conclusion by an unrelated approach, we investigated whether binding to known target mRNAs is influenced by ANKS3 or ANKS6 or vice versa. Bicc1 binds ANKS3 through SAM:SAM interactions and on Ank repeats, independently of KH domains (Rothé et al., 2018). Here, yeast-two-hybrid and ANSK3 pull-down assays identified the Bicc1 KH domain repeat as a novel interface. Since we could not identify a unique ANKS3 region that mediates this novel interaction with the KH repeat, we predict that it consists of several epitopes that act additively or redundantly to limit access of RNA to one or several KH domains. Our finding that ANKS3 inhibited Bicc1 binding to its own mRNA and to Dand5 and Adcy6 3’-UTRs strongly supports this model. ANKS3 also competed with the Dand5 3’- UTR for GST-Bicc1 fusions in cell-free assays. Conversely, the restoration of Bicc1 condensation by ANKS6 overexpression correlated with increased RNA binding. Interestingly, ANKS3 was also outcompeted by RNA when we only mutated its SAM:SAM interface with Bicc1, instead of adding ANKS6. These findings suggest that ANKS proteins regulate mRNA binding of Bicc1 through specific multivalent interactions that govern the topology of its heterooligomerization to either promote liquid-liquid or gel-like phase transitioning (**Figure 7**).

Recently, CLIP-seq analysis in in HEK293T cells identified >1500 mRNAs that can bind to overexpressed Bicc1 (Estrada Mallarino et al., 2020). Endogenous *BICC1* and *DAND5* are not transcribed in those cells, and ADCY6 mRNA levels are exceedingly low. However, both our gain-of-function studies in HEK293T cells and the regulation of endogenous Bicc1 by Anks3 knockdown in mIMCD3 cells support the conclusion of our cell-free assays that the threshold of mRNA binding is greatly increased by Anks3-mediated inhibition, reducing the background noise of non-specific binding and allowing ANKS6 to spatiotemporally couple the recruitment of specific target mRNAs to Bicc1 phase transitioning in metronomically regulated amounts. While Dand5 mRNA is also below detection in murine IMCD3 kidney cells, its 3’-UTR has been validated *in vivo* as a conserved target mediating left-right axis specification in response to cilia-induced activation of Bicc1 preferentially on the future left side (Maerker et al., 2021; Minegishi et al., 2021). Here, we observed increased binding of endogenous Bicc1 to its own mRNA and to endogenous *Adcy6* transcripts upon RNAi depletion of ANKS3 in IMCD3 cells. Interestingly, this increase was accompanied by an unexpected corresponding increase in Bicc1 mRNA itself, but not of its protein, consistent with a possible conserved role for Bicc1 in regulating its own expression by negative feedback (Chicoine et al., 2007). By contrast, Adcy6 mRNA expression remained unchanged. These observations warrant future studies to investigate whether ANKS3 selectively reduces the stability of Bicc1 mRNA, for example to influence the access of other client RNAs, and whether Bicc1 autoregulation is physiologically relevant.

The novel dual inhibition of Bicc1 by ANKS3 and of ANKS3 by ANKS6 agrees with the known functions of these proteins during development (Rothé et al., 2020). In addition, ANKS6 is implicated in a program of reversible renal tubule dilation that allows urine flow to flush out calcium oxalate crystals during adult tissue homeostasis (Torres et al., 2019b, 2019a). An ANKS6 point mutation in the SAM domain that inhibits this program and promotes cystic growth inhibits the association with ANKS3 (Bakey et al., 2015; Leettola et al., 2014). Protein network stiffening and transition of specific RNA-binding proteins towards near-solid states is a frequent pathological consequence of specific mutations, for example in amyotrophic lateral sclerosis or frontotemporal dementia patients (Portz et al., 2021). Therefore, and since spontaneous mutations in Bicc1 and ANKS6 mostly affect their SAM domains (Bakey et al., 2015; Cogswell et al., 2003; Flaherty et al., 1995; Kraus et al., 2012; Nauta et al., 1993; Neudecker et al., 2010), future studies could address whether the dynamics of this protein network during phase transitioning are generally perturbed in ciliopathies.

## METHODS

### Antibodies

Monoclonal rabbit anti-HA (Sigma H6908), monoclonal mouse anti-FLAG® M2 (Sigma F3165), polyclonal rabbit anti-ANKS6 (Sigma HPA008355), polyclonal rabbit anti-CNOT1 (LSBio LS-C335614), polyclonal rabbit anti-GAPDH (Abcam ab70699) and monoclonal mouse anti- yTubulin (Sigma T6557) antibodies were used for Western-blot analyses. Immunoprecipitation and detection of endogenous Bicc1 in IMCD3 cells were performed using custom-made affinity-purified polyclonal rabbit anti-Bicc1 antibody (Rothé et al., 2015).

### Plasmids

The plasmids pEF-SA::Dand5-3’UTR (Minegishi et al., 2021), pCMV-SPORT6::HA-Bicc1, ΔKH and ΔSAM have been described previously (Maisonneuve et al., 2009). The plasmid pGEX- 1λT::Bicc1-KH has been previously described (Piazzon et al., 2012). PCR fragments of individual KH domains and full-length Bicc1 WT or mutD have been amplified from appropriate pCMV- SPORT6 constructs (Rothé et al., 2015) and cloned between BamHI and XhoI sites of pGEX-1λT and pGEX-6p1, respectively. To derive pCMV-SPORT6::HA-Bicc1 ΔKH-mutD, a BstBI-XbaI fragment of pCMV-SPORT6::HA-Bicc1 mutD (Rothé et al., 2015) was used to replace the corresponding fragment in pCMV-SPORT6::HA-Bicc1 ΔKH. Plasmid pCMV-GFP::Bicc1 was constructed by cloning the AgeI-XbaI fragment of pCMV-SPORT6::HA-Bicc1 into XmaI-XbaI sites of pCMV-EGFP-C1(Neo). The ΔKH derivative has been obtained by replacing the internal BsrGI/XmaI fragment encoding Pro_41_-Gly_451_ by a synthetic linker. Alternatively, to express GFP- Bicc1 without HA tags, Bicc1 cDNA including its short 5’UTR was fused to GFP in pCMV-GFP-C1 as an AgeI-XbaI fragment derived from the Bicc1 IMAGE clone 2655954 (Genbank AF319464). To generate doxycycline-inducible TRE3G-eGFP-Bicc1-puro, HA-tagged GFP-Bicc1 from pCMV-GFP::Bicc1 was cloned as an AgeI-XbaI fragment into XmaI-XbaI of a derivative of pLenti CMVTRE3G Puro DEST (w811-1) lacking the Gateway cloning site (provided by David Suter, EPFL, together with lentiviral plasmid hPGK-rtTA3G-IRES-bsd.

Plasmid pGBKT::ANKS3 and fusions of human BICC1 FL (full-length) cDNA, KH repeat, IVS or SAM domain with the activation domain of GAL4 in pACT2 have been described (Rothé et al., 2018). PCR amplicons of the KH_1_, KH_2_ and KHL (which encompasses KHL1, KH3 and KHL1 domains) were digested with BglII and XhoI and inserted between BamHI and XhoI sites of pACT2.

To generate pCS-Luc::Dand5-3’UTR, a fragment encompassing the 3’end of the CDS (from nucleotide 539) and the 3’UTR of *Dand5* was amplified by PCR and inserted between the XhoI and BglII sites of pCS-Luc-link-MS2. To increase the number of MS2 tandem repeats, a synthetic 12xMS2 cassette (GeneArt Gene Synthesis, ThermoFisher) was cloned into XhoI/HpaI-digested pCS-Luc-link-MS2 to derive pCS-Luc::12xMS2. Insertion of 12xMS2 into BglII-SnaBI of pLuc::Dand5-3’UTR and into the XbaI site of pCMV-GFP::Bicc1 gave rise to pCS-Luc::Dand5- 3’UTR-12xMS2 and pCMV-GFP::Bicc1-12xMS2, respectively. UbiC::NLS-HA-2xMCP-HALO was from Addgene (Halstead et al., 2015).

### Cell lines and cell culture

IMCD3 (CRL-2123), HEK293T (CRL-11268) and HeLa cell lines (CCL-2) were purchased from ATCC and cultured in DMEM (Sigma) supplemented with 10% FBS (Sigma), 1% GlutaMAX (Thermo Fisher Scientific), and 1% gentamicin (Thermo Fisher Scientific). IMCD3.sgBicc1; tetO::GFP-Bicc1; rtTA cells were derived from the CRISPR/Cas9-edited IMCD3.sgBicc1 clone #14 (Leal-Esteban et al., 2018) by stable transduction with hPGK-rtTA3G-IRES-bsd and then, in clonal cell lines, with TRE3G-eGFP-Bicc1-puro lentivirus. To induce GFP-Bicc1 and analyze its condensation in the cytoplasm or in cell extracts, IMCD3.sgBicc1;tetO::Bicc1; rtTA cells at a density of 2×10^6^ per 10 cm dish were treated with 0.5 μg/mL doxycycline for 24 hrs, or as indicated. To assess effects on GFP-Bicc1 mobility, cells were treated with blebbistatin (SelleckChem, 50uM for 30min), cytochalasin D (Sigma, 0.5 µM for 1h), monastrol (SelleckChem, 100 µM for 12h), Y27632 (SelleckChem, 10 µM for 2h) or nocodazole (Sigma, 100 ng/ml for 12h). To deplete ANKS3, IMCD3-sh::ANKS3 cells (Schlimpert et al., 2018) were seeded at a density of 2×10^6^ per 10 cm dish and treated with 0.25 μg/mL doxycycline. After two days, cells were passaged into two 15 cm dishes at a density of 5×10^6^ cells per plate and doxycycline was added in fresh complete medium every 48 hrs for 5 days.

### In vitro transcription

The cDNA templates were provided with the SP6 RNA polymerase promoter and with the sequence tag TGTCTGGGCAACAGGCTCAGG at their 5’ and 3’ ends, respectively, using Overlap Extension PCR primers and Phire Green Hot Start II PCR Master Mix (ThermoFisher). After agarose gel electrophoresis, PCR amplicons were purified on NucleoSpin® Gel and PCR Clean- up (Macherey-Nagel). Template DNAs of interest (500 ng each) were transcribed *in vitro* during 2 hours at 37°C using SP6 RNA polymerase kit (Roche), followed by a 1 hr treatment with DNAse I (Roche) prior to purification of the newly synthesize RNA on Quick Spin Columns (Roche) and quantification by OD_600_ measurement.

### Purification of GST fusion proteins

Fusions of glutathione S-transferase (GST) with Bicc1 fragments in plasmid pGEX-1λT or with full-length Bicc1 in plasmid pGEX-6p1 were expressed in *E. coli* BL21 (Novagen) as described (Piazzon et al., 2012) and purified using glutathione-Sepharose 4B (GE Healthcare). Purification was carried out in a buffer consisting of 50 mM Tris-HCl, pH 8, 200 mM NaCl, 1 mM dithiothreitol (DTT).

### Sedimentation of membraneless compartments by stepwise differential centrifugation

HEK293T were transfected with 2 μg HA-Bicc1 or HA-Bicc1 ΔKH, or 4 μg HA-Bicc1 ΔSAM per 10 cm dish using jetPEI (Polyplus Transfection) 1 d before the experiment. Transfected HEK293T cells, or stable IMCD3.sgBicc1; tetO::GFP-Bicc1; rtTA cells (one 10 cm dish per condition) were washed with ice-cold PBS and passed eight times through a syringe needle (no. 30) in extraction buffer containing 50 mM Tris-HCl (pH 7.6), 5 mM MgCl_2_, 50 mM NaCl, 1 mM dithiothreitol (DTT), 0.1% Nonidet P-40 (NP-40), RNasin (Promega), phosphatase inhibitors (Sigma), and protease inhibitors (Roche). After a first centrifugation at 2,000 g for 2 min at 4°C, the extracts were centrifuged by steps of increasing speed from 4,000 g to 10,000 g for 5 min at 4°C each. After gentle collection of the supernatants, the pellets were resuspended after each step in the same volume of extraction buffer as the initial cell pellet. To assess whether GFP-Bicc1 sedimentation is salt-sensitive, aliquots of the supernatant of the 2,000 g centrifugation step were supplemented with increasing concentrations of NaCl and incubated for 20 min at 4°C before sequential centrifugation at 4,000 g and 10,000 g. Alternatively, to assess whether the sedimentation in heavy fractions depends on RNA, supernatants from the 2,000g centrifugation step were incubated for 20 min at RT with 0.4 μg/μL of RNase A or with 0.3 μg/μL of *in vitro*- transcribed RNA prior to centrifugation at 4000 g and 10,000 g. For Western blot analysis of heavy and light fractions, 15 µl aliquots of each were dissolved in Laemmli buffer, size-separated on reducing SDS PAGE gels, transferred to nitrocellulose membranes, blocked with skim milk and incubated over night with the indicated antibodies, and with fluorescently labelled secondary antibodies for analysis on a Odyssey CLx scanner (LI-COR Biosciences). To image GFP-Bicc1, heavy or light fractions were each mixed with equal volumes of 4% low melting agarose at 37°C, and mounted under coverslips. Pictures were acquired on a Zeiss LSM700 confocal microscope.

### Tracking of GFP Bicc1 granules

After treatment with 1 µg/mL doxycycline for 24 hrs on 35 mm glass bottom plates (Ibidi), IMCD3.sgBicc1; tetO::GFP-Bicc1; rtTA cells were placed in PBS and imaged at 37°C using a TCS SP8 STED 3x microscope (Leica Microsystems) equipped with white-light laser. Images were acquired using a 63x objective with a 7.5x optical zoom and pinhole size of 5 Airy Units (AU). Image format and laser speed were set at 512 x 512 pixels and 1400, respectively. Images were taken every 15 seconds intervals for 15 min along the Z-axis. Time-lapse videos were generated after Z-projection (maximum intensities) of all images. Dynamic parameters were obtained using ImageJ TrackMate plugin and R statistical software. The parameters to extract Links, Spots and Tracks statistics from Trackmate were as follows: Estimated blob diameter, 0.6 µm; Filter spots, “Mean intensity”; Linking Max Distance, 2 µm; Gap-closing max distance, 1 µm; Gap-closing max frame gap, 0 µm. For a given recording, individual spot displacement values were normalized based on the Track length, which corresponds to the number of consecutive images where a spot was successfully tracked.

### FRAP analysis

FRAP analysis in HeLa cells was performed using a Zeiss LSM710 confocal microsocope. ROI were photobleached with maximum laser power at 488 nm or 561 nm wavelengths. Images were taken during 4 min at 2.5 sec intervals with a fully open pinhole. Results were expressed as the percentage of recovery relative to the baseline of residual fluorescence intensity after photobleaching at t = 0 sec, and normalized to the fluorescence of an unbleached area. The half-time of recovery was estimated by linear regression.

### Structured Illumination Microscopy

HeLa cells on cover slips with a thickness of 170 μm were transfected with 1 μg of plasmid per well in a 6-well plate. After 24 hrs, cells were fixed in methanol for 10 min at −20°C and imaged on a 3D NSIM Nikon microscope using an objective APO TIRF100x/1.49 oil combined with an additional lens 2.5x. GFP-Bicc1 was detected using a Coherent Sapphire 488 nm laser (200 mW) and imaged using an Andor iXon3 897 camera. The acquisition time was 100 ms at a readout of 3 MHz. 3D images were reconstructed from stacks of 15 images (5 phase shifts and 3 rotations) with a z-step of 120 nm. Pictures of the final 3D volume rendering were generated using Imaris software.

### Immunofluorescence staining

HeLa cells transfected with 1 μg of plasmid per well in 6-well plates were incubated and one day later passed to sterile coverslips in 24-well plates. After incubation for another day, cells were fixed in methanol for 10 min at −20°C. After washing in PBS, coverslips were incubated 1 hr at RT in blocking buffer containing PBS and 1% BSA, and 2 hrs at RT in blocking buffer containing anti-HA antibody (Sigma). After rinsing with PBS, cells were incubated with anti-rabbit Alexa 647 in blocking buffer for 1 hr at RT in presence of DAPI (1/10000). Pictures were acquired on a Zeiss LSM700 confocal microscope.

### Protein-RNA colocalization by time-lapse imaging

3×10^5^ HeLa cells in 35 mm glass bottom dishes (Ibidi) were transfected with 2xMCP-NLS-HaloTag together with GFP-Bicc1-12xMS2 (1 µg each), or, where indicated, with GFP-Bicc1 and either Empty- or Dand-3’UTR-12xMS2 reporters. After 36 hrs, cells were incubated for 15 min with the HaloTag® TMR G8252 ligand (Promega) in fresh medium. Thereafter, unbound TMR was removed by washing 3 times with culture medium (including a final incubated incubation in fresh medium for 30 min) and 3 times in PBS. Imaging was performed in PBS supplemented with 2% BSA on an inverted Zeiss LSM700 confocal microscope. Four stacks were acquired every 1.5 min during at least 15 min. Movies were mounted with a speed of 5 pictures/sec.

### Protein co-immunoprecipitation assay in HEK293T cells

HEK293T cells were transfected with 2 µg GFP-Bicc1, HA-Bicc1 or HA-Bicc1 ΔKH per 10 cm dish, or with 4 µg HA-Bicc1 mutD or HA-Bicc1 ΔKH-mutD. 24 hrs later, cells from two dishes per condition were washed with ice-cold PBS and pooled in extraction buffer containing 20 mM Tris-HCl (pH 7.4), 2.5 mM MgCl_2_, 100 mM NaCl, 5% glycerol, 1 mM dithiothreitol (DTT), 0.05% Nonidet P-40 (NP-40), RNasin (Promega), phosphatase inhibitors (Sigma), and protease inhibitors (Roche). Total cell extracts were prepared by ultrasonication followed by two rounds of centrifugation at 10,000 × g for 5 min at 4°C to remove debris. Supernatants were incubated with anti-HA beads (Sigma) for 2 hrs at 4°C on a rotating wheel. After washing four times for 10 min in 20 mM Tris-HCl pH 7.4, 2 mM MgCl_2_, 200 mM NaCl, 1 mM DTT, and 0.1% NP-40, beads were suspended in Laemmli buffer and loaded on reducing SDS PAGE gels to size-separate eluted proteins. For immunoblot analysis, proteins were transferred to nitrocellulose membranes, blocked with skim milk and incubated over night with the indicated antibodies, and with fluorescently labelled secondary antibodies for analysis on a Odyssey CLx scanner (LI-COR Biosciences).

### RNA co-immunoprecipitation assay in IMCD3 and HEK293T cells

HEK293T cells were transfected with 2 μg *Dand5*-3’UTR, HA-Bicc1 and ANKS3-Flag plasmids each per dish, and 8 μg v5-ANKS6. HEK293T cells from two 10 cm dishes, or IMCD3- sh::ANKS3 cells from two 15 cm dishes per condition, were washed with ice-cold PBS, extracted with 20 mM Tris-HCl (pH 7.4), 2.5 mM MgCl_2_, 100 mM NaCl, 5% glycerol, 1 mM dithiothreitol (DTT), 0.05% Nonidet P-40 (NP-40), RNasin (Promega), phosphatase inhibitors (Sigma), and protease inhibitors (Roche), by passing them eight times through a syringe needle (no. 30). Extracts were centrifuged twice at 10,000 × g for 5 min at 4°C. and 5% of each set aside as controls for the “input”. For immunoprecipition, 20 μL of Protein G-sepharose beads (GE Healthcare) coated with 12 µL rabbit anti-Bicc1, or 20 µL of mouse anti-HA beads (Sigma) and preabsorbed with RNAse-free BSA (800 µg/mL) were incubated with the remainder of each extract for 2 hrs at 4°C on a rotating wheel, then rinsed four times 10 min in wash buffer (20 mM Tris-HCl pH 7.4, 2 mM MgCl_2_, 200 mM NaCl, 1 mM DTT, and 0.1% NP-40). While 10% of the beads were analyzed by immunoblotting as described above, the remainder and half of each input sample were subjected to phenol-chloroform extraction. After ethanol precipitation, isolated RNA was treated with RQ1 DNase (Promega) and converted to cDNA by PrimeScript Reverse Transcriptase Kit (Takara). The resulting cDNA was subjected to PCR or qPCR analysis using Phire Green Hot Start II PCR Master Mix (Promega) or GoTaq qPCR Master Mix (Promega), respectively, using the primers For- GCTGAGCATCCTAGAGGAATGC and Rev-TAAACCCATGACTGGGGGACCATGTCTAG for the 3’-UTR of Dand5 mRNA, or For-ACAGAGCCTCGCCTTTGCC and Rev-CTCCATGCCCAGGAAGGAAGG for β-actin mRNA. The amount of co-immunoprecipitated mRNA as a percentage of the input was calculated with the formula: 100 × 2[(Ct(Input) − log2(100/2.5) − Ct(IP)], where the value 2.5 represents 2.5% of the original cytoplasmic extract, and where Ct is the cycle threshold for qPCRs on input or immunoprecipitate (IP) samples. Fold enrichment was calculated relative to cells transfected with the corresponding empty vector for HA-Bicc1 and normalized relative to the amount of HA-Bicc1 bait in the IP fraction.

### Reconstitution of multiprotein complexes and RNP by GST pull-down

To reconstitute RNPs, a fluorescent DNA probe 5’-CTGAGCCTGTTGCCCAGAC-3’ carrying a 5’-Dynomics 681 dye (Microsynth AG) was pre-annealed to the complementary 3’-tag of *in vitro*-transcribed Dand5-3’UTR_66-110_ or Dand5-3’UTR_226-270_ RNAs by denaturation for 3 min at 98°C and renaturation for 10 min at RT. Subsequently, the labelled RNA (100 pmol) was incubated with GST-KH fusion protein during 1 hr at 4°C on wheel. To assess binding of preassembled RNPs or of GST-Bicc1 fusion proteins alone to ANKS proteins, HEK293T cells were transfected with ANKS3-Flag and v5-ANKS6 and extracted as described above. Cleared extracts were incubated for 2 hrs at 4°C with glutathione-Sepharose 4B beads that were coated with GST alone (control), or with GST-Bicc1 fusions or RNPs. Proteins that were bound to the beads were washed and analyzed by Western blotting as described above. Retention of the RNA was monitored using an Odyssey CLx Infrared Imaging System (LI-COR Biosciences) by imaging the annealed fluorescent probe directly in the gel before immunoblotting of associated proteins. Binding of the GST fusions was validated by Coomassie staining of eluted proteins. GST alone was included in all experiments as a specificity control, but after prolonged migration that was required to resolve the protein of interest at the top of the gel could not be retained due to its small size.

### Yeast two-hybrid assay

Binding of ANKS3 to various domains of human BICC1 was assessed in reciprocal yeast two-hybrid assays as described (Rothé et al., 2014) by fusing each as a bait to the DNA-binding domain of the GAL4 transcription factor (GAL4-BD) in plasmid pGBKT7 (Clontech), and as a prey to the activation domain of the GAL4 transcription factor (GAL4-AD) in plasmid pACT2 (Clontech). To monitor the induction of a HIS3 reporter gene by complexes of bait and prey fusion proteins, appropriate pairs of pACT2 (LEU2) and pGBKT7 (TRP1) plasmids were transformed into haploid *S. cerevisiae* CG1945 cells (mat a; ura3-52, his3-200, ade2-101, lys2-801, trp1-901, leu2-3, 112, gal4-542, gal80-538, cyhr2, LYS2::GAL1UAS-GAL1TATA-HIS3, URA3::GAL417-mers(x3)- CYC1TATALacZ) and strain Y187 (mat α; gal4, gal80, ade2-101, his3-200, leu2-3,112, lys2-801, trp1-901, ura3-52, URA3::Gal1UAS GAL1TATA-LacZ). Diploid progeny from crossings on YPD medium and selected on Leu^-^, Trp^-^ medium were plated on Leu^-^, Trp^-^, His^-^ medium for 3 days at 30°C to select the cells where reconstituted GAL4 AD/BD complexes induced the HIS3 reporter gene. Where indicated, 3-Amino-1, 2, 4-triazol (3-AT) was added as a competitive inhibitor of histidine synthesis to evaluate the strength of the interactions.

## ACKNOWLEDGEMENTS

The authors would like to thank Dr. Gerd Walz for kindly providing the IMCD3-sh::ANKS3 cell line, and Dr. Soeren Lienkamp for v5-ANKS6 plasmid. We thank Dr. Bruno Charpentier (Université Lorraine-Nancy) for comments on the manuscript. This work was supported in part by the Human Frontiers Science Program fellowship LT000216/2016 to S.F., and by Rare Diseases GRS-051/13 grant from Gebert Rüf Stiftung to D.B.C, and resources and services of the Bioimaging Research Core Facility and the Gene Expression Core Facility at the School of Life Sciences of EPFL.

## AUTHOR CONTRIBUTIONS

Conceptualization and Methodology, B.R., S.F. and D.B.C.; Validation, B.R. and D.B.C.; Formal analysis and Investigation, B.R. and S.F; Data Curation, B.R.; Writing – Original Draft, B.R. and D.B.C.; Writing – Review & Editing, B.R., S.F. and D.B.C.; Visualization, B.R.; Supervision, D.B.C.; Project Administration, D.B.C.; Funding Acquisition, S.F. and D.B.C.

## SUPPLEMENTARY FIGURE LEGENDS

**Figure 1 - figure supplement 1.**
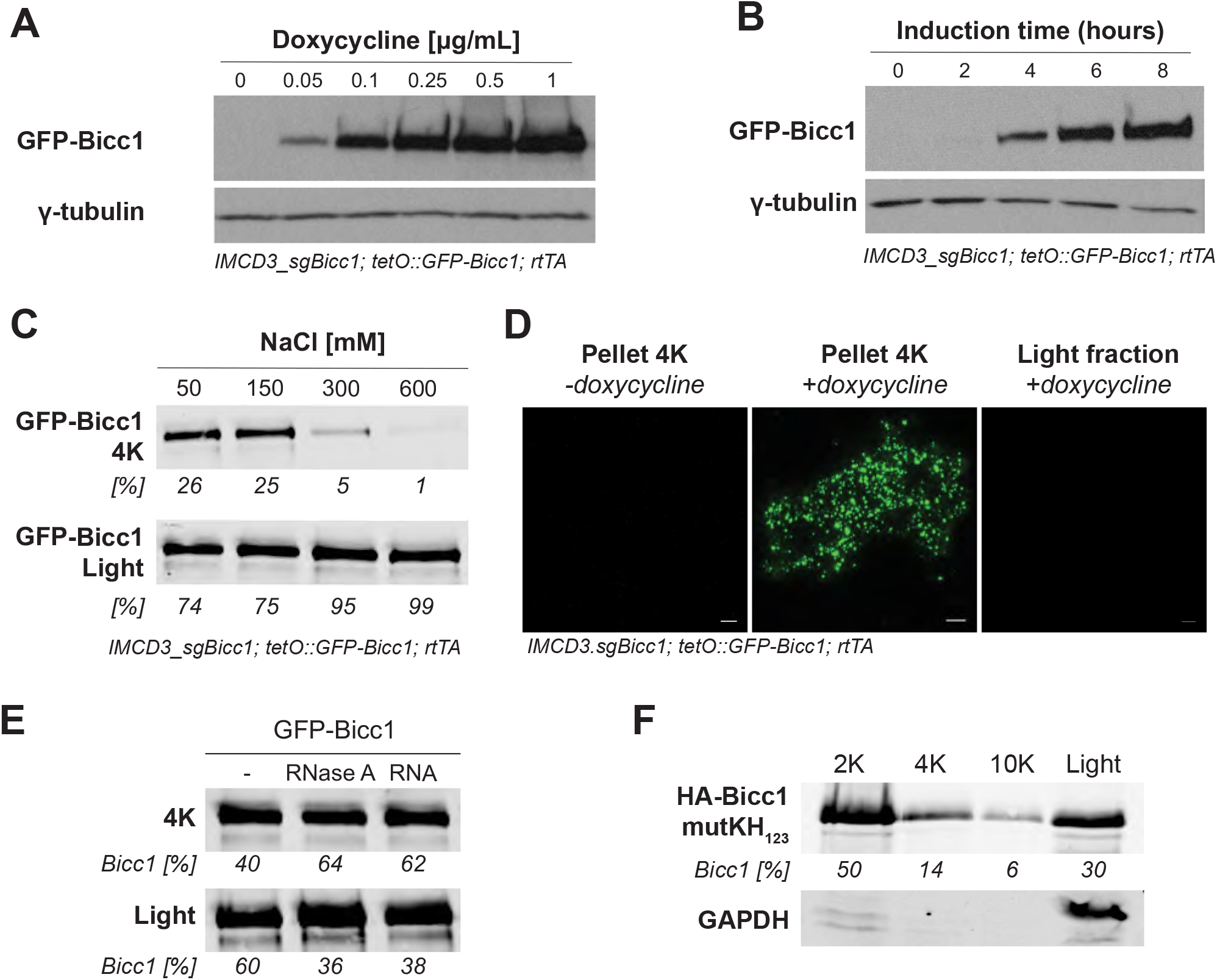
Inducible expression and confocal imaging of GFP-Bicc1 condensates in cell extracts, and their solubilization in high salt. **(A, B)** Western blots of sgBicc1 mutant IMCD3 cells treated with increasing concentrations of doxycycline (A) and for the indicated durations (B) to monitor the induction of a tetO::GFP-Bicc1 transgene by stably transduced reverse Tet transactivator (rtTA). **(C)** Distribution of GFP-Bicc1 between the 4K and light fractions in the presence of increasing concentrations of NaCl. **(D)** Confocal microscopy of GFP-Bicc1 in the indicated light and heavy (4K) fractions. (**E**) Distribution of GFP-Bicc1 between the 4K and the Light fractions isolated by differential centrifugation of inducible IMCD3 cells. Before centrifugation, total lysates were treated with RNAse A or supplemented with *in vitro* transcribed AC6-3’UTRprox RNA. **(F)** Sedimentation of HA-Bicc1 mutKH_1,2,3_ by differential centrifugation. GAPDH was used as a marker of the cytoplasmic light fraction. Data are means + SD from at least four independent experiments. *p < 0.05, **p < 0.01, ***p < 0.001 (Student’s t-test).

**Figure 1 - figure supplement 2.**
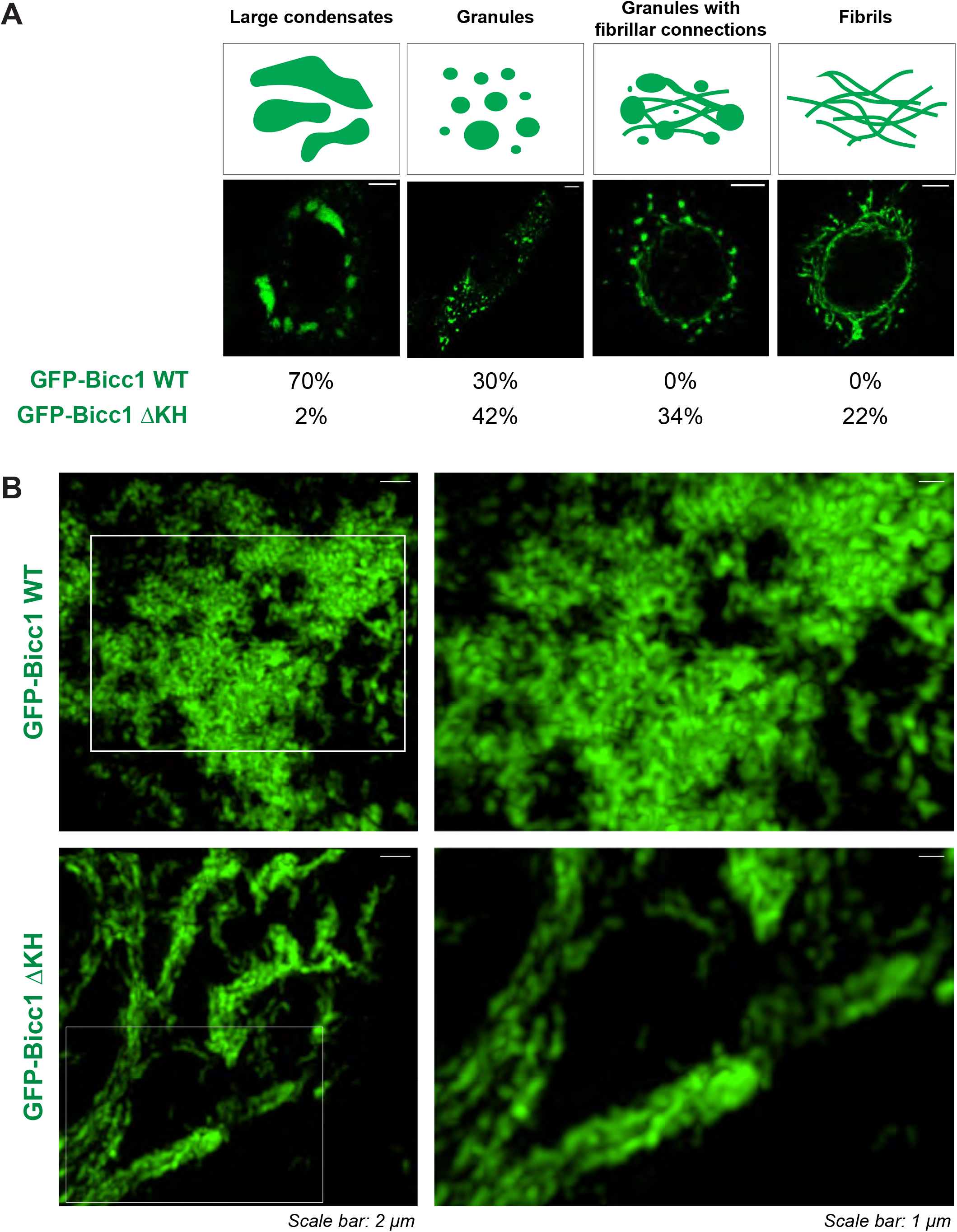
Internal topology of GFP-Bicc1 granules by high resolution microscopy. **(A)** Relative frequencies of the GFP-Bicc1 distribution patterns observed by confocal imaging in transiently transfected HeLa cells (n=125). Four classes were defined base on the size and the morphology of the GFP-Bicc1 condensates: individual entities with size < 3 µm were defined as “Granules”, whereas larger entities were defined as “Large condensates”. GFP-Bicc1 ΔKH specific patterns showing fibrillar structures were classified as “Granules with fibrillar connections” or “Fibrils” depending the presence or the absence of round-shape granules. **(B)** Representative 3D SIM images of GFP-Bicc1 WT and ΔKH transiently expressed in HeLa cells and corresponding to “Large condensates” and “Fibrils” patterns, respectively.

**Figure 2 - figure supplement 1.**
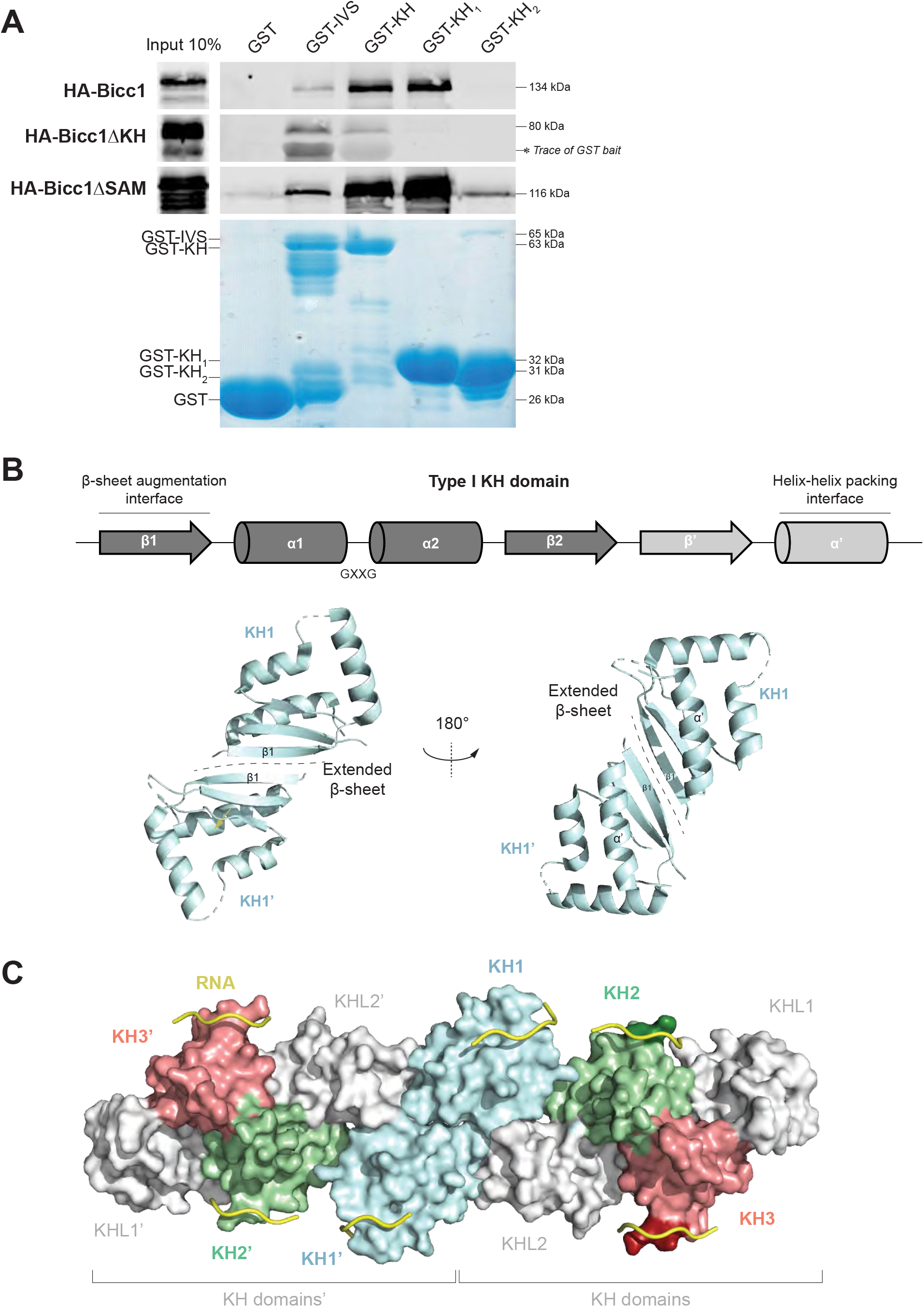
Model of a Bicc1 KH dimer. **(A)** Pull-down of HA-Bicc1 WT, ΔKH and ΔSAM from HEK293T cell extracts by glutathion sepharose beads coated with recombinant domains of Bicc1 fused to GST. **(B)** Secondary structure organization of a type I KH domain (top) and X-ray structure of the hBICC1 KH_1_ dimer (PDB: 6GY4) in cartoon representation (below). Structure elements conserved at the interface between other KH domains are indicated. **(C)** Dimerization of the Bicc1 KH tandem repeat via KH_1_ domains, modeled *in silico* based on the X-ray structure in (B). The RNA was docked in the RNA-binding grooves by alignment with the solution structure of the third KH domain of KSRP bound to a G-rich target sequence (PBD: 4B8T) using the PyMol program (Schrödinger). Docking was done manually without further steps of energy minimization.

**Figure 3 - figure supplement 1.**
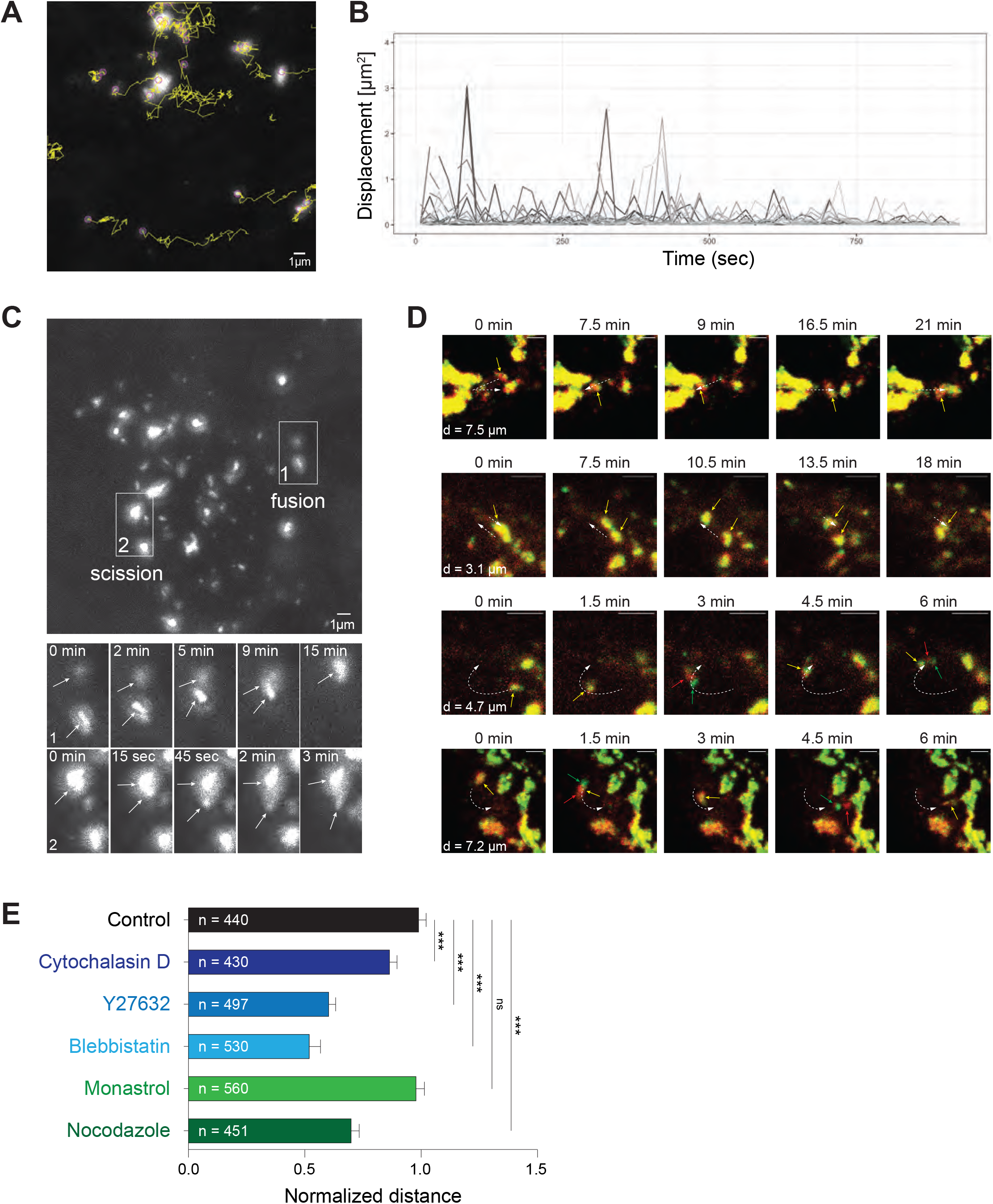
Actomyosin-dependent mobility of GFP-Bicc1 granules and reversible liquid-gel phase separation of bound mRNAs. **(A)** Displacement tracks (yellow) of inducible cytoplasmic GFP-Bicc1 granules in IMCD3 cells. **(B)** Quantification of the Squared Displacement of GFP-Bicc1 foci. The threshold of 1 µm^2^ was used to define non-Brownian motion. Events of granules leaving the focal plane are indicated by interrupted lines. **(C)** Time-lapse imaging of fusion and scission events of GFP-Bicc1 granules. **(D)** Time-lapse imaging of RNPs of GFP-Bicc1 (green) with its own 12xMS-tagged mRNA (red) in HeLa cells. White arrows represent trajectories of directional movements. Covered distances (d) are given on the left. Scale bars, 2 µm. **(E)** Normalized distance travelled by GFP-Bicc1 granules in IMCD3 cells before and after treatment with the indicated inhibitors. Movements were tracked as in (D). Data are means + SD from n individual granules. ***p < 0.001 (Student’s t-test).

**Figure 5 - figure supplement 1.**
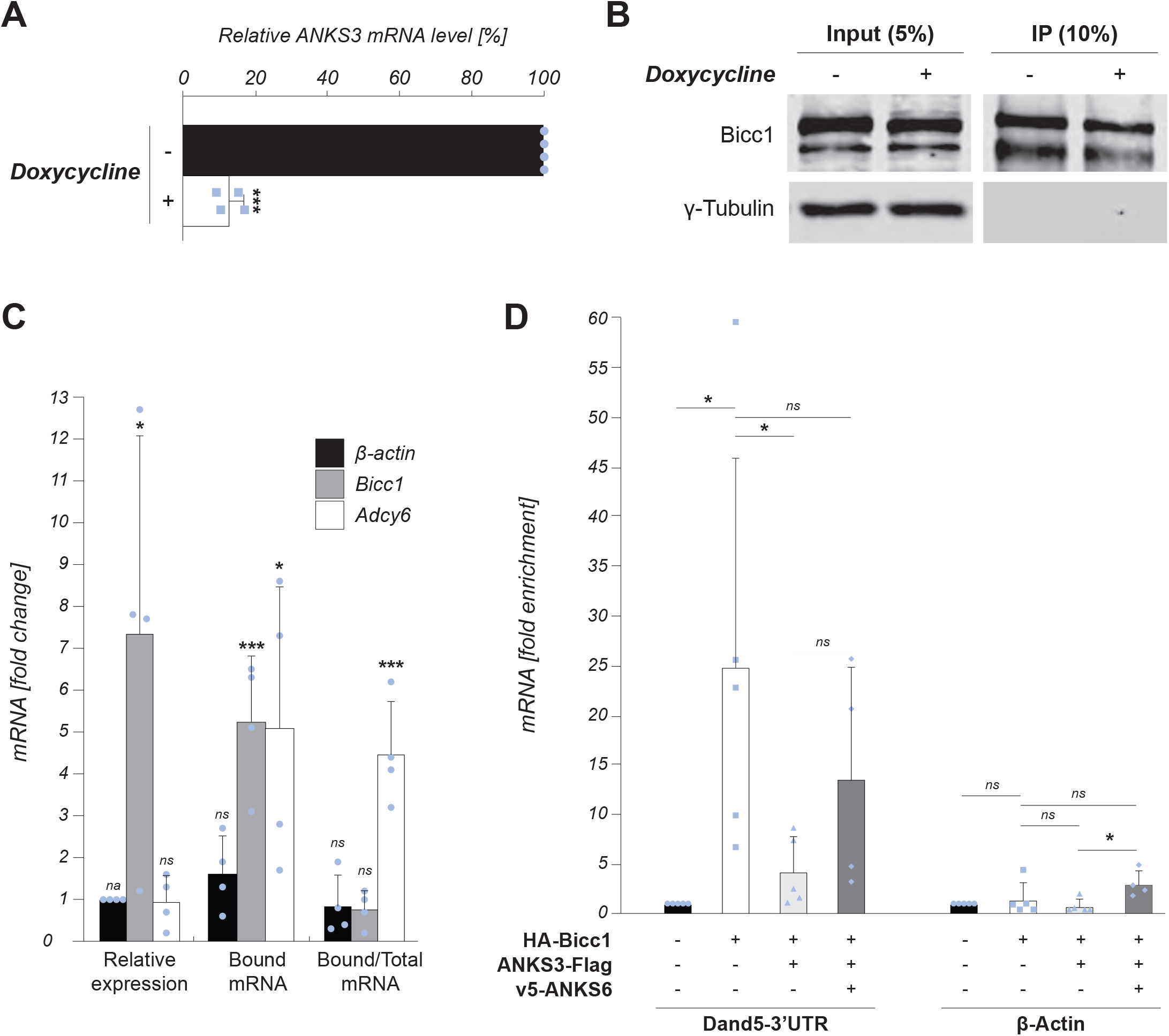
ANKS3 stimulates the RNA binding activity of endogenous Bicc1 in IMCD3 cells. **(A)** RT-qPCR analysis of endogenous *ANKS3* mRNA in IMCD3 cells treated with or without 0.25 µg/ml doxycycline (Dox) to induce the expression of Anks3 shRNA. The *ANKS3* transcript level relative to the β-actin mRNA signal is expressed as a percentage of the baseline in untreated cells. **(B)** Western blot of endogenous Bicc1 in cytoplasmic extracts (inputs) and immunoprecipitates of IMCD3 cells treated with or without doxycycline to induce *ANKS3* shRNA. γ-Tubulin was a loading control. **(C)** RT-qPCR analysis of the indicated mRNAs in IMCD3 before and after ANKS3 depletion in IMCD3 cells relative to β-actin. For each category, the values are expressed as the fold change normalized to the untreated condition. The category “Bicc1-bound” represents the amount of mRNA found in Bicc1 immunoprecipitates from ANKS3-depleted cells normalized to the Bicc1 protein signal in the IP fraction, and expressed relative to the control without ANKS3 depletion. “Bound/Total mRNA” represents the ratio of mRNA in Bicc1 immunoprecipitates relative to the mRNA in the input. β-actin mRNA is the negative control used to define the background of non-specific RNA binding. Data are means + SD from four independent experiments. ns: non-significant, na: non-applicable, *p < 0.05, **p < 0.01, ***p < 0.001 (Student’s t-test). **(D)** Additional analysis of the RT-qPCR data shown in Figure 5A. Ratios of co-immunoprecipitated mRNAs over the input, normalized to the amount of HA-Bicc1 in the IP fraction and then expressed relative to the corresponding condition without HA-Bicc1. β-actin mRNA served as a negative control to visualize the level of unspecific RNA binding by Bicc1. Data are means + SD from at least four independent experiments. ns: non-significant, *p < 0.05, **p < 0.01, ***p < 0.001 (Student’s t-test).

**Figure 6 - figure supplement 1.**
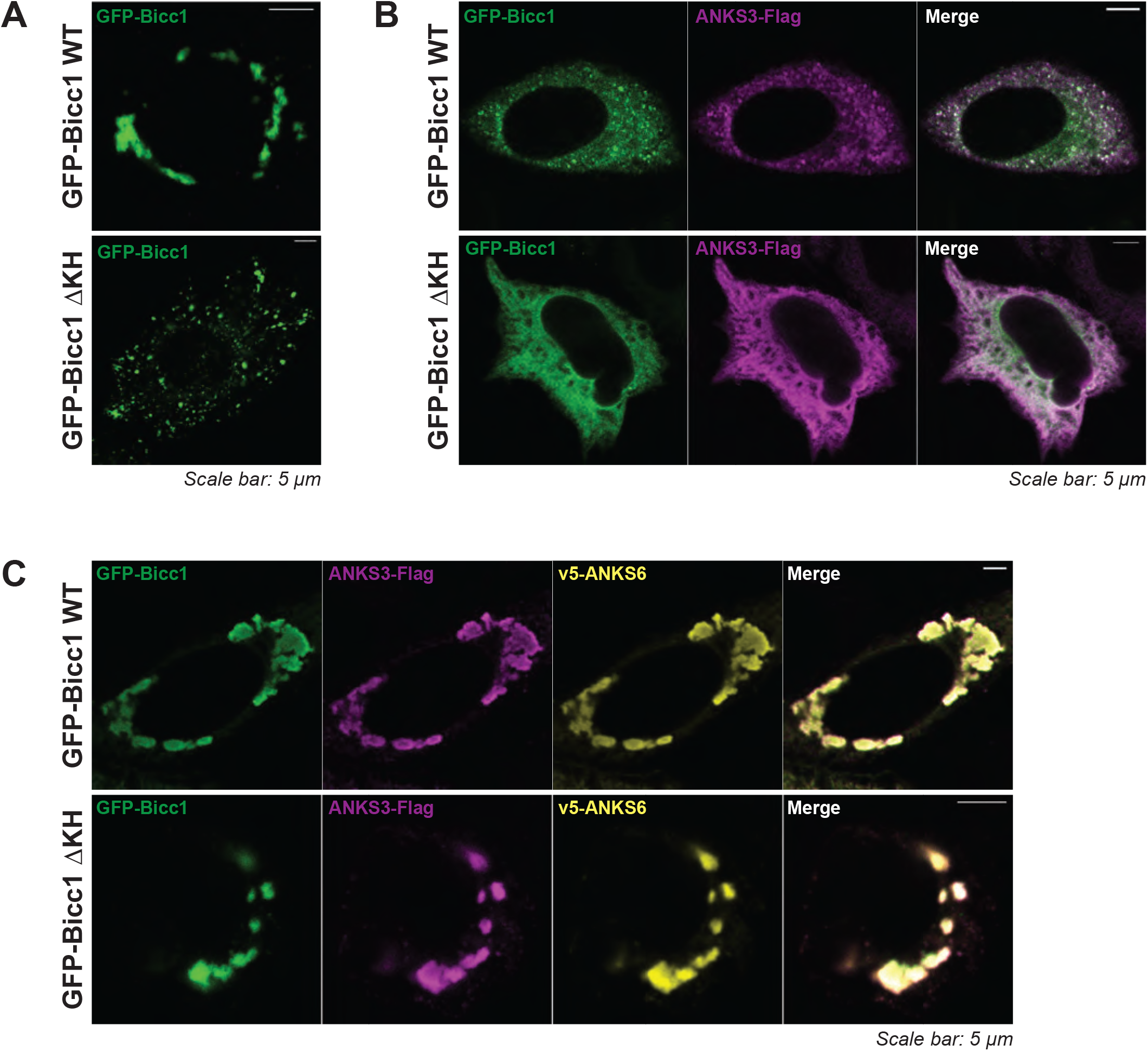
Cytoplasmic clustering of GFP-Bicc1 WT and ΔKH is inhibited by ANKS3 and rescued by ANKS6. **(A-C)** HeLa cells transiently transfected with GFP-Bicc1 WT or ΔKH alone (A), or co-transfected with ANKS3-Flag (B), or ANKS3-Flag and v5-ANKS6 (C). ANKS3-Flag and v5-ANKS6 were stained by immunofluorescence using anti-Flag and anti-ANKS6 antibodies, respectively.

## SUPPLEMENTAL MOVIES

**Figure 3 - video 1-2.** Liquid-gel phase transitioning of GFP-Bicc1 protein (green) and its 12xMS2- tagged mRNA (red) imaged in real time in two representative HeLa cells during 15 min, starting 24 hours after transfection.

**Figure 3 - video 3.** Time-lapse imaging of GFP-Bicc1-12xMS2 mRNA (red) entering a large GFP- Bicc1 protein condensate (green) upon their collision.

**Figure 3 - video 4.** Time-lapse imaging of GFP-Bicc1-12xMS2 mRNA (red) entering a small GFP- Bicc1 protein droplet (green), followed by fusion of the resulting RNP particle (yellow) with a massive pre-existing Bicc1 condensate.

**Figure 3 - video 5.** Time-lapse imaging of a GFP-Bicc1 RNP condensate (yellow) where the protein (green) dissociates from its 12xMS-tagged mRNA (red) during the course of approximately 10 min.

**Figure 6 - video 1-3.** Time-lapse imaging of GFP-Bicc1 (green) and its own 12xMS2-tagged mRNA (red) in HeLa cells. GFP-Bicc1-12xMS2 was expressed alone (Figure 6 - video 1), or in combination with ANKS3-Flag (Figure 6 - video 2), or ANKS3-Flag and v5-ANKS6 (Figure 6 - video 3). ANKS3- Flag unmixes small GFP-Bicc1 foci from mRNA, whereas ANKS6 rescues large GFP-Bicc1 condensates and restores their stable co-localization with GFP-Bicc1 mRNA.

